# Within-population genome size variation is mediated by multiple genomic elements that segregate independently during meiosis

**DOI:** 10.1101/623470

**Authors:** Claus-Peter Stelzer, Maria Pichler, Peter Stadler, Anita Hatheuer, Simone Riss

**Author notes:** Corresponding Author University of Innsbruck, Research Department for Limnology, Mondseestr. 9, 5310 Mondsee, Austria.

## Abstract

Within-species variation in genome size has been documented in many animals and plants. Despite its importance for understanding eukaryotic genome diversity, there is only sparse knowledge about how individual-level processes mediate genome size variation in populations. Here we study a natural population of the rotifer *Brachionus asplanchnoidis* whose members differ up to 1.9-fold in genome size, but were still able to interbreed and produce viable offspring. We show that genome size is highly heritable and can be artificially selected up or down, but not beyond a minimum diploid genome size. Analyses of segregation patterns in haploid males reveal that large genomic elements (several megabases in size) provide the substrate of genome size variation. These elements, and their segregation patterns, explain the generation of new genome size variants, the short-term evolutionary potential of genome size change in populations, and some seemingly paradoxical patterns, like an increase in genome size variation among highly inbred lines. Our study suggests that a conceptual model involving only two variables, (1) a minimum genome size of the population, and (2) a vector containing information on additional elements that may increase genome size in this population (size, number, and meiotic segregation behaviour), can effectively address most scenarios of short-term evolutionary change of genome size in a population.

## Introduction

Despite an exponential increase of genomic information during the last two decades, there is still no consensus about the ultimate causes of genome size variation in eukaryotes [1-3]. At the core of this controversy is the puzzling genome size variation across eukaryotic taxa, spanning approximately five orders of magnitude. Genome sequencing has revealed that this variation is primarily caused by gene-, chromosome-, or genome duplications, by variation in the length of introns, number of transposons, and the amount of simple repetitive DNA [4-7]. Since most of these sequences make no substantial contribution to the phenotype, at least not through their information content, genome size is only a poor predictor of organismal complexity [1]. On the other hand, ubiquitous correlations of genome size and cell size, or other phenotypic traits such as metabolic- or developmental rates, and body size [8-12], suggest the sheer amount of DNA in a genome can affect the phenotype[13]. Collectively, this might explain why current theories on genome size variation in eukaryotes differ strongly in their emphasis on selection, mutation and drift [2, 3, 13-16].

Theories invoking selection for increased genome size have been criticized for assuming causal links behind the correlations between genome size and phenotypic traits [2]. Indeed, such correlations often involve species that have been separated for long evolutionary timespans and thus differ in many other aspects than genome size. According to the mutational hazard hypothesis, non-coding DNA is never beneficial, but it may accumulate as a consequence of genetic drift [17]. Thus, this hypothesis offers an alternative, neutral explanation to the observed genome size - phenotype correlations by stating that the accumulation of non-coding DNA in organisms with large body size might be due to their smaller effective population sizes [17, 18]. Testing whether large genome size can sometimes be beneficial, or whether it is at least conditionally deleterious, ideally requires a model system that exhibits substantial genome size differences across a relatively homogeneous genomic background. This requirement appears to be best fulfilled in species with intraspecific genome size variation, where individuals share their genomic background and evolutionary history.

Cases of intraspecific genome size variation are well-documented in plants [19], with cultivated maize and its close relatives being probably one of the best-studied examples [20, 21]. In animals, intraspecific genome size variation has been found in snapping shrimp [22] and in grasshoppers [23]. Interestingly, two intensively studied model species with comparably small genomes, *Arabidopsis thaliana* and *Drosophila melanogaster*, have also turned out to exhibit substantial levels of intraspecific genome size variation [24, 25], suggesting that this phenomenon might be more widespread than previously assumed. Intraspecific genome size variation is sometimes associated with variation in chromosome numbers, for instance due to supernumerary (B-)chromosomes [19], but there are also documented cases where genome size variation is not reflected the karyotype [22, 26].

Despite its importance, surprisingly little is known about the basic mechanisms and inheritance of intraspecific genome size variation, and the links to population-level phenomena, i.e.: What characterizes those parts of a genome that account for the differences in genome size between individuals? How is genome size inherited by offspring? How (fast) can the trait ‘genome size’ change through generations, e.g., if it is directionally selected? Even in the best-studied systems, researchers typically rely on assumptions and draw analogies to models of quantitative genetic variation. For example, the trait ‘genome size’ is often considered a quantitative trait influenced by a large number ‘loci’, with ‘alleles’ that increase or decrease genome size [27, 28]. We are not aware of any empirical study that has yet assessed the appropriateness of such a model.

Here we study the basic mechanisms of genome size variation in a population of the rotifer *Brachionus asplanchnoidis*. This species is characterized by an almost two times larger genome size relative to its sister species, by a high 44% genomic content of repetitive elements [29], and by intraspecific genome size variation [30, 31]. Using crossbreeding experiments, selfed lines, and artificial selection, we disentangle the basic mechanism by which variation for genome size is mediated in this population, and how it is inherited by offspring. We capitalize on several advantages of our model system, such as short generation times, sexual and asexual reproduction, and a haploid-diploid lifecycle, which allows us to probe into meiotic patterns associated with intraspecific genome size variation. We could identify the size and number of individual elements that contribute to increases in genome size, and thus account for the gradual differences among individuals in the population. Our findings suggest that intraspecific genome size variation can be conceptualized in terms of a ‘minimum genome size’ (representing the smallest genomes in a population), and additional ‘elements’ found in individuals with larger genomes (each characterized by size, number, and meiotic segregation behaviour). This difference in perspective, compared to a conventional quantitative trait-model, has significant implications on the genome size distribution of populations, and the short-term evolutionary potential of the trait ‘genome size’.

## Results

### Within-population genome size variation in *B. asplanchnoidis*

To quantify genome size variation within populations, we examined 118 *B. asplanchnoidis* clones sampled from four geographic populations (Fig. 1, Supplementary table 1). Of these, two Austrian populations from ‘Obere Halbjochlacke’ (OHJ, 74 clones) and ‘Runde Lacke’ (RL, 29 clones) show highly significant intrapopulation variation (OHJ: ANOVA F_52,194_ = 112, *P* < 0.001; RL: F_28,84_ = 23.14, *P* < 0.001). The OHJ-population spans a genome size range of 1.33-fold, from 414 to 552 Mbp (Fig. 1a), with one outlier at 792 Mbp (i.e., 1.91-fold). The RL-population is also variable (Fig. 1b), spanning a range of 1.24-fold across all sampled clones. In contrast, the Lake Nakuru population (11 clones sampled) was not significantly variable (F_10,21_ = 8.64, *P* = 0.078), and most genomes were close to 420 Mbp (Fig 1c). Previously, we reported two conspecific clones isolated from a Mongolian lake, which have relatively large genome sizes of 652 and 732 Mbp (Fig. 1c, data from Riss et al. 2016). Genome size is mitotically stable, since the genome sizes of clones, as well as the differences among clones, were highly reproducible over a period of more than five years (i.e., approximately 600 asexual generations, Supplementary Fig. 1).

**Fig. 1.**
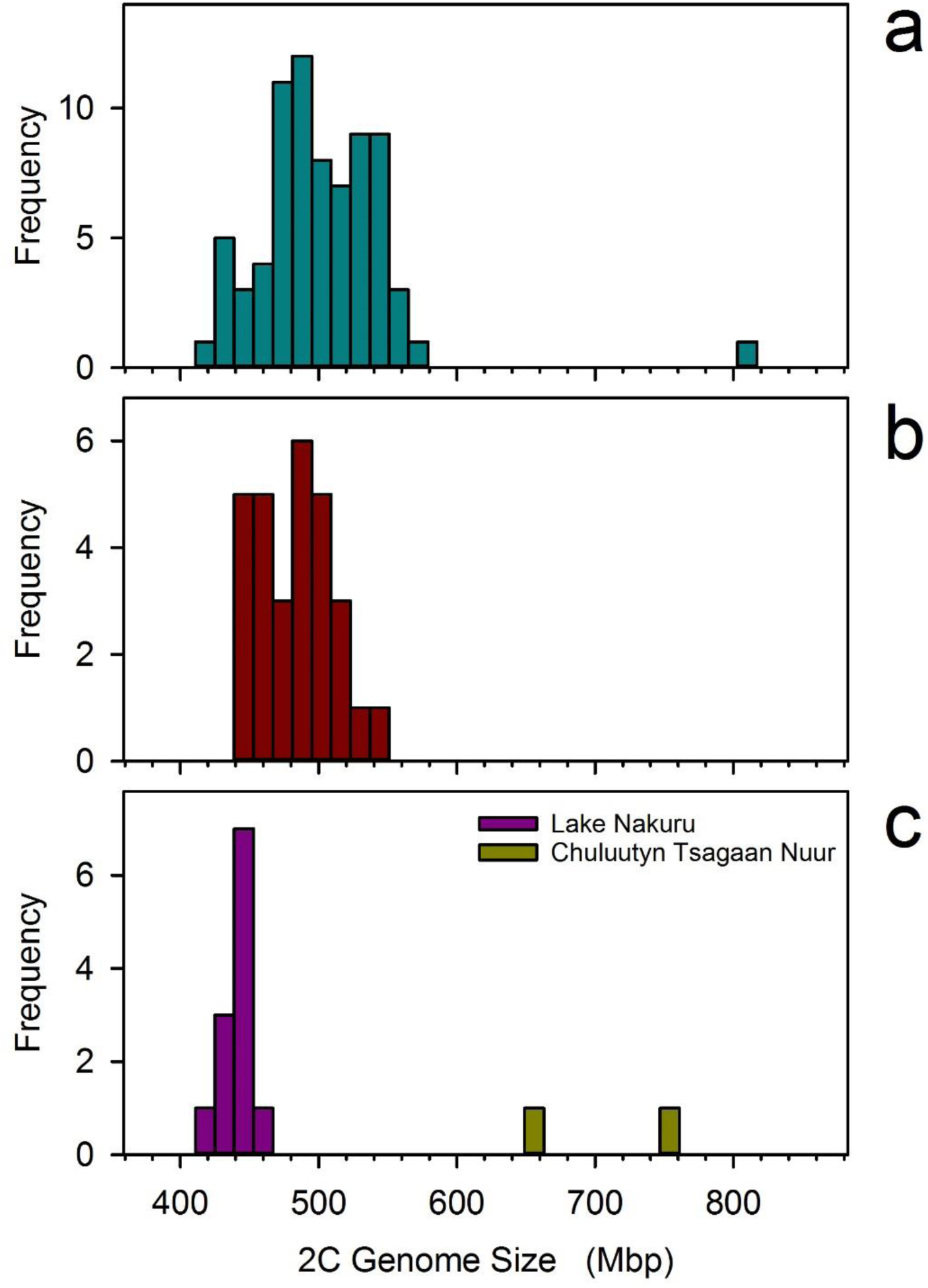
Genome size variation in four natural *B. asplanchnoidis* populations. **a** Obere Halbjochlacke (Austria). **b** Runde Lacke (Austria). **c** Lake Nakuru (Kenya) and Chuluutyn Tsagaan Nuur (Mongolia). Genome size was highly variable in the two Austrian populations (genome sizes ranging from 414 to 792 Mbp), whereas clones isolated from Lake Nakuru sediments differed little from each other (426 ± 5.1 Mbp, mean and standard deviation). The two clones from the Mongolian site had distinct and relatively large genome sizes of 652 and 732 Mbp (Data from [32]).

### Inheritance of within-population genome size variation

Clones with divergent genome size can mate with each other and produce viable and fertile offspring, which are intermediate in genome size between their parents but show some variation (Supplementary Fig. 4, see also [32]). Genome size responds to artificial selection with extremely high heritability. We applied truncation selection to the OHJ population by crossing clones with the 10% largest genome sizes among each other (excluding the outlier clone at 792Mbp), and by crossing clones with the 10% smallest genomes, respectively. In the up-selection treatment, we obtained genome sizes exceeding the range of the parental OHJ population (Fig. 2). We could select genome sizes of up to 640 Mbp, with a narrow-sense heritability *h*^2^ of 0.905 in the first generation, and 0.912 in the second generation. Likewise, heritability was high in the first generation of the down-selection treatment (h^2^ = 0.924). However, in contrast to selection for large genome size, it did not extend the range of the original population. In fact, we could not select genome sizes below 414 Mbp. Additionally, h^2^ of the second generation of the down-selection treatment collapsed to zero. A parent-offspring regression including all our crosses yields an overall estimate for h^2^ of 0.96 (Supplementary Fig. 5).

**Fig. 2.**
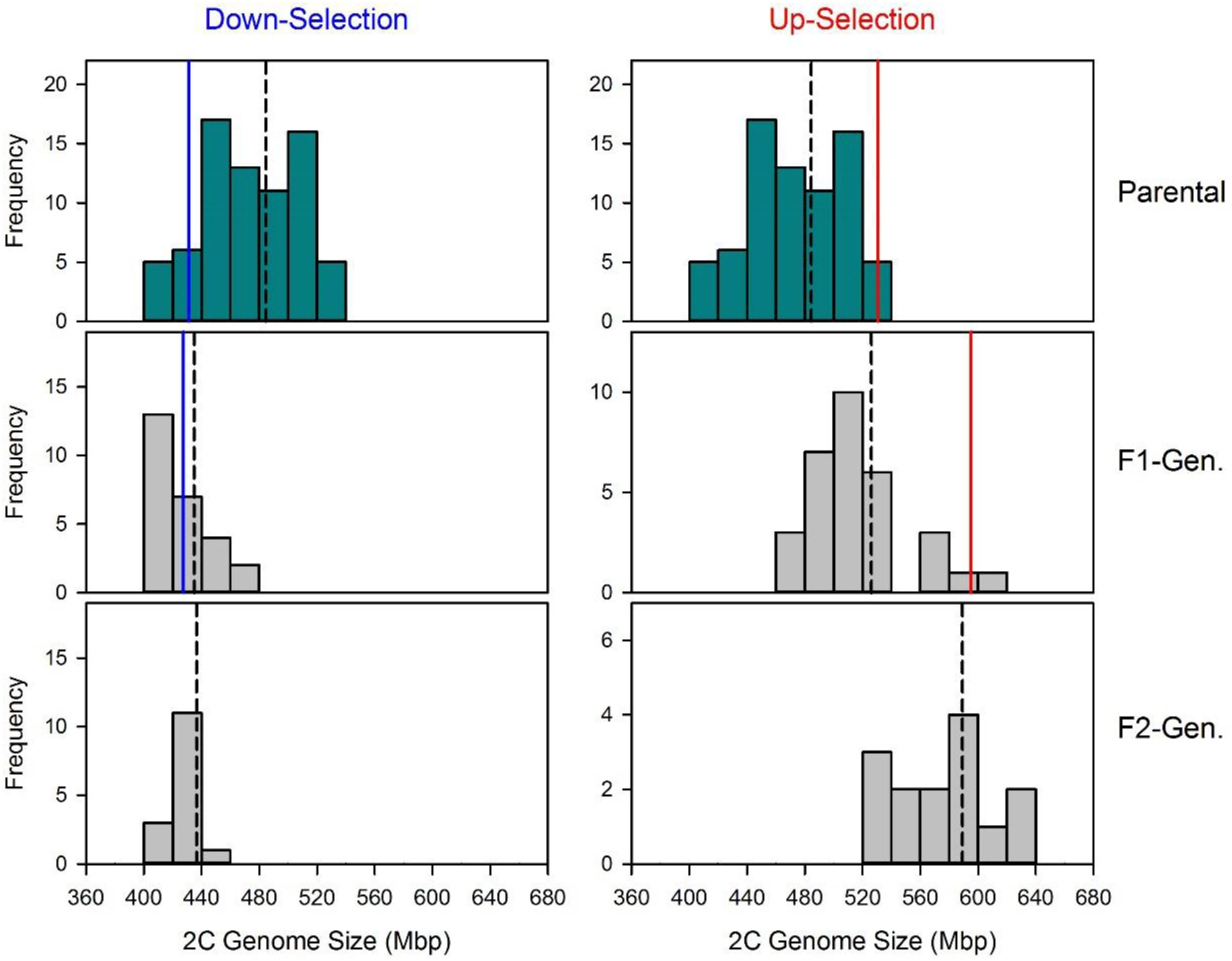
Artificial selection for increased or reduced genome size. The parental population consisted of clones isolated from the natural population from Obere Halbjochlacke (cf., Fig. 1a) and was the same for both treatments. Dashed black lines indicate population mean genome sizes for each generation. Solid lines indicate the mean genome size of the selected top 10% (in red) or bottom 10% (in blue) for the parental and F1-generation.

Combining all available genome size estimates of *B. asplanchnoidis* reveals a positively skewed distribution, with a high number of observations at 410-430 Mbp and an elongated tail of large genome sizes (Supplementary Fig. 6). The striking absence of genome sizes smaller than 410 Mbp, despite high sampling effort and intentional selection for small genome size, suggests that this genome size might be a true biological limit. By contrast, there was no constraint in terms of increases in genome size, as suggested by the more or less continuous rise of genome sizes to 792 Mbp. Interestingly, this clone with the largest genome size was not artificially selected, but hatched from a resting egg of the natural population (c.f., Fig. 1a).

To get additional insights into the inheritance of genome size, we analysed two selfed lines that were derived from a clone with large and small genome size, respectively. Theoretically, one would expect trait variation to decrease upon selfing as this causes 50% reduction of heterozygosity each generation. In contrast, we found that genome size variation remained high in a selfed line that descended from a large-sized clone (524 Mbp). Genome sizes of selfed offspring ranged between values of 522 Mbp and 644 Mbp, representing a 23% increase (Fig. 3). Among-clone variation in the large selfed line was statistically significant, even after three generations of selfing (ANOVA *F*_9,24_ = 53.61, *P* < 0.001). Variation in the large line was also significantly higher than in the line descending from the small genome (*p* < 0.001; R package *cvequality*, version 0.2.0, [33]). After three generations of selfing, we performed a sexual cross between both lines. As was the case in the cross between two natural clones, the cross of the selfed lines yielded offspring that were variable and intermediate between their parents (Fig. 3).

**Fig. 3.**
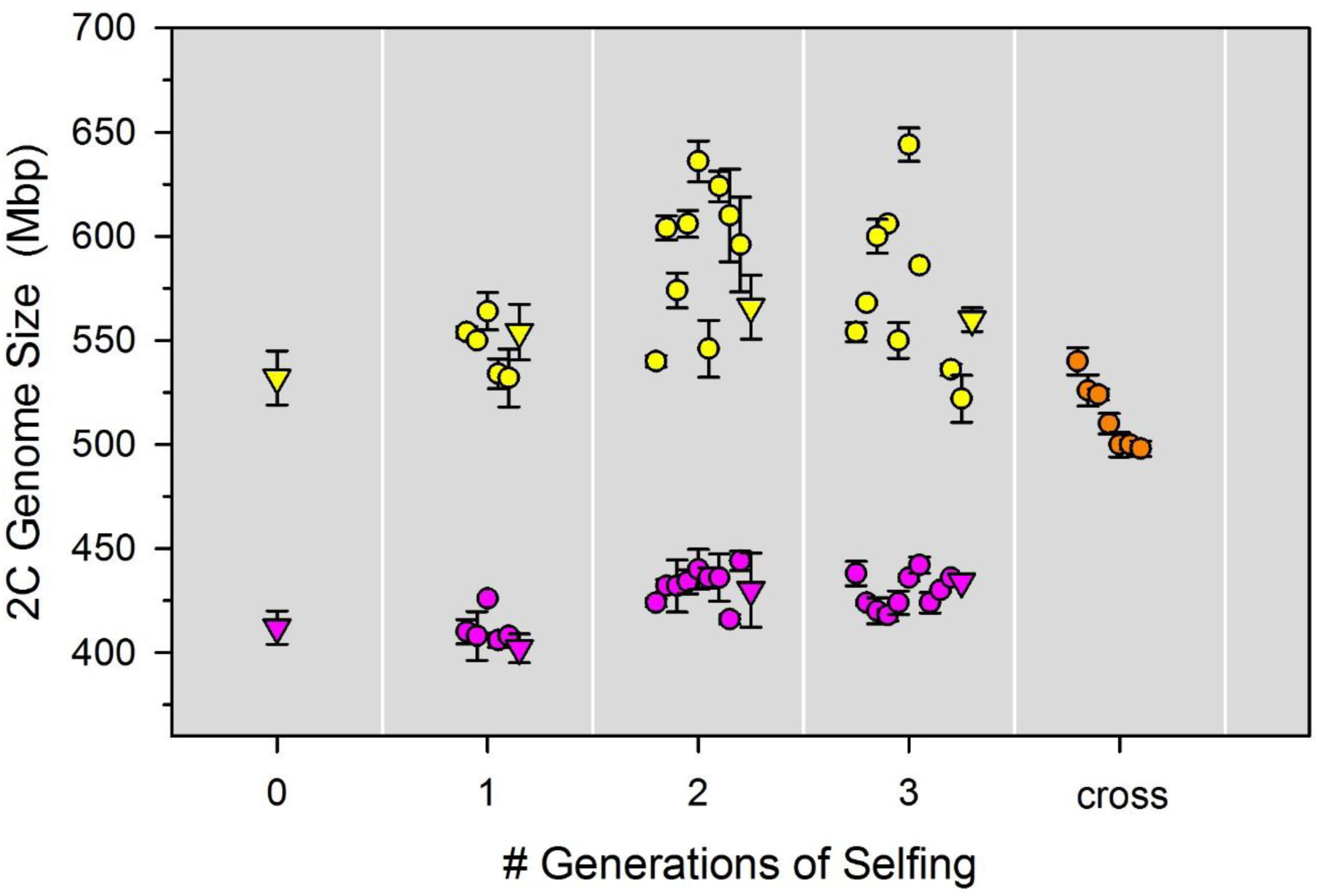
Genome size variation in two selfed lines and their inter-line cross. Two selfed lines were established from two natural rotifer clones, one with large genome size (524 Mbp, yellow) and one with small genome size (414 Mbp, purple), and were propagated for three sexual generations. Clones used for breeding are indicated by triangles, their siblings are displayed as circles. After three generations of selfing, the two lines were crossed with each other to produce hybrid offspring (orange).

### Evidence for independently segregating genomic elements

To gain mechanistic insights into segregation of genome size variation during meiosis, we analysed haploid rotifer males. In a diploid organism with equally sized chromosome pairs and no extra-chromosomal elements, all meiotic products should contain exactly half the DNA of a diploid cell. Consistently, haploid males of the rotifer *B. calyciflorus* have half the genome size of females ([34]; see Appendix S2 in this publication). We also find this pattern in some of our *B. asplanchnoidis* clones (e.g. Fig. 4a). However, in many others we obtained striking variation in male genome size, which manifested in multiple discrete “male peaks” (Fig. 4b-h, Supplementary table 2). In the simplest case, we observed two male peaks spaced symmetrically around the expected 1C-value (Fig. 4b). In general, the male peak pattern of a clone could be characterized by (i) the number of peaks, (ii) an odd/even number of peaks, indicating the presence/absence of a central male peak at exactly 1C, and (iii) the relative abundance of certain male genome size classes, as inferred from the area under each male peak. In most cases, the central male genome sizes (the ones close to 1C) were more abundant than the ones in the periphery (Supplementary Fig. 7, Supplementary table 2). Overall, male peak patterns were clone-specific and highly repeatable (Supplementary Fig. 8).

**Fig. 4.**
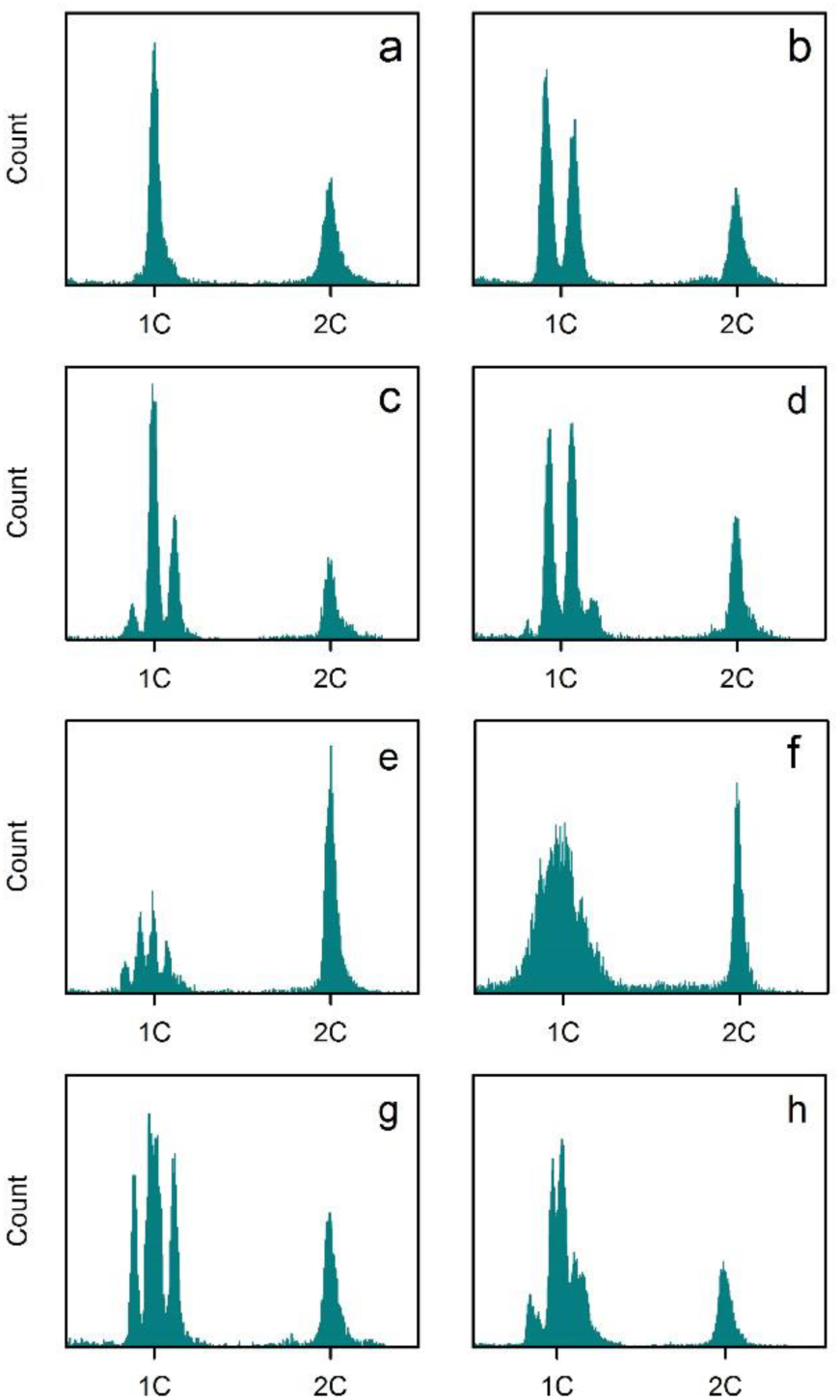
Representative examples of genome size variation in haploid males. Each histogram corresponds to a different clone. X-axis corresponds to the YL2-A fluorescence on a linear scale. Only the section surrounding the 1C and 2C values is shown. The 1C value was set at 0.5x the median fluorescence of the 2C (female) peak. **a** One single “male peak” (MP), indicates that male genome size is not variable. Note that the MP is located at exactly one half (=1C) of the diploid female genome size (2C). **b** Two MPs located at 0.46 and 0.54 of female genome size, indicating that there are two discrete classes of male genome size. **c** Three MPs, located at 0.44, 0.5, and 0.56. **d** Four MPs, located at 0.41, 0.47, 0.53, and 0.59. **e** Five MPs, located at 0.42, 0.46, 0.5, 0.54, and 0.58 (the MP on the far right is very faint). **f** No discrete MPs, but extremely broad distribution of male genome sizes around the 1C value. However, the 2C peak of this sample is very narrow (∼2% CV), indicating that the broad MP is due to real variation in genome size rather than a lack of precision. **g** Four MPs. Note that in contrast to **d**, the MPs are approximately in the same height (i.e., relative frequency), and that there is unequal spacing between the MPs (the middle MPs are close together). **h** Six MPs with unequal spacing, showing some resemblance to the case of three MPs, but each MP appears to be split up into two sub-peaks.

In some clones, we could not resolve individual male peaks, but instead obtained an extremely broad distribution of genome sizes around the 1C value (e.g., Fig 4f). This pattern likely reflects real genome size variation among males of a clone, rather than a lack of measurement precision, since the coefficient of variation of male genome sizes in this example was 12%, while variation of the female peak ∼2%. Interestingly, this clone with the extremely broad male peak was the same as the “outlier” with the largest genome size of 792 Mbp (Fig. 1a). Although this clone was the extreme case, we obtained broad distributions of male genome sizes in other clones as well (data not shown). Finally, in some clones we observed characteristically unequal distances between male peaks (Fig. 4 g,h), such that some male peaks were more narrow together than others.

All the subsequent results and analyses build on the following *ad hoc* hypothesis, which is based on our observations so far: We hypothesized that large genomic elements, several Mbp in size, are segregating independently from each other, and thus give rise to discrete classes of male genome size. By independent, we mean that each element has an approximately equal chance of segregating into any of the four gametes during gametogenesis. Note that this is different from normal chromosome segregation, where homologous chromosomes will always end up in opposite gametes. The presence of one independently segregating element can explain the simplest pattern of two male peaks (Fig 4b), being present in the large male genome size but absent in the small one. Similarly, two equally sized elements can explain a pattern of three male peaks (Fig. 4c), which correspond to genome size classes with zero, one, or two elements. Likewise, patterns with higher numbers can be explained by *n*-1 elements. Assuming that independently segregating elements have identical size, we can predict the relative ratios of the male genome size classes to 1:1 (one element), 1:2:1 (two elements), or 1:3:3:1 (three elements). Indeed, some of our clones closely follow these predicted frequencies, whereas others showed some deviations (Supplementary fig. 7). For example, in clones with two male peaks, the smaller male genome size was often at higher frequency than the expected value of 0.5 (Supplementary fig. 7), suggesting that males without an element were more frequent among the hatched males. Likewise, in clones with three and four male peaks, the central male genome sizes were sometimes at a higher frequency than expected. One clone showing four male peaks with almost identical heights (Supplementary fig. 7c) presents a particularly interesting deviation. In this same clone, the two central peaks were closer together (cf. Fig. 4g). The most parsimonious explanation seems to be that this clone carries two differentially sized elements, a small and a large one, and that the four male peaks correspond to: (1) zero elements, (2) the small element, (3) the large element, and (4) both elements. Likewise, two large and one small element can explain the “three double-peaks” pattern in the clone depicted in Fig. 4h.

With this in mind, we estimated the size and number of independently segregating elements, based on the distance between male peaks in a clone and its 2C genome size (Fig. 5a). We find that the natural OHJ-population harbours a large diversity of elements (Fig. 5b). Many clones contain elements of ∼34Mbp size, but smaller elements down to 15 Mbp were also present in this population. In contrast, the clones of our “large” selfed line apparently contained only elements in the 34 Mbp size range (Fig. 5b), and they exhibited significantly less variation in element size than the natural population (p<0.001; R package *cvequality*, version 0.2.0, [32]). Interestingly, the founding clone of this line shows four male peaks (Fig. 4d), and indicated three elements of ∼34Mbp size. Thus, it appears that this same 34Mbp element causes all the observed genome size variation in the “large” selfed line, being present in different numbers in different offspring.

**Fig. 5.**
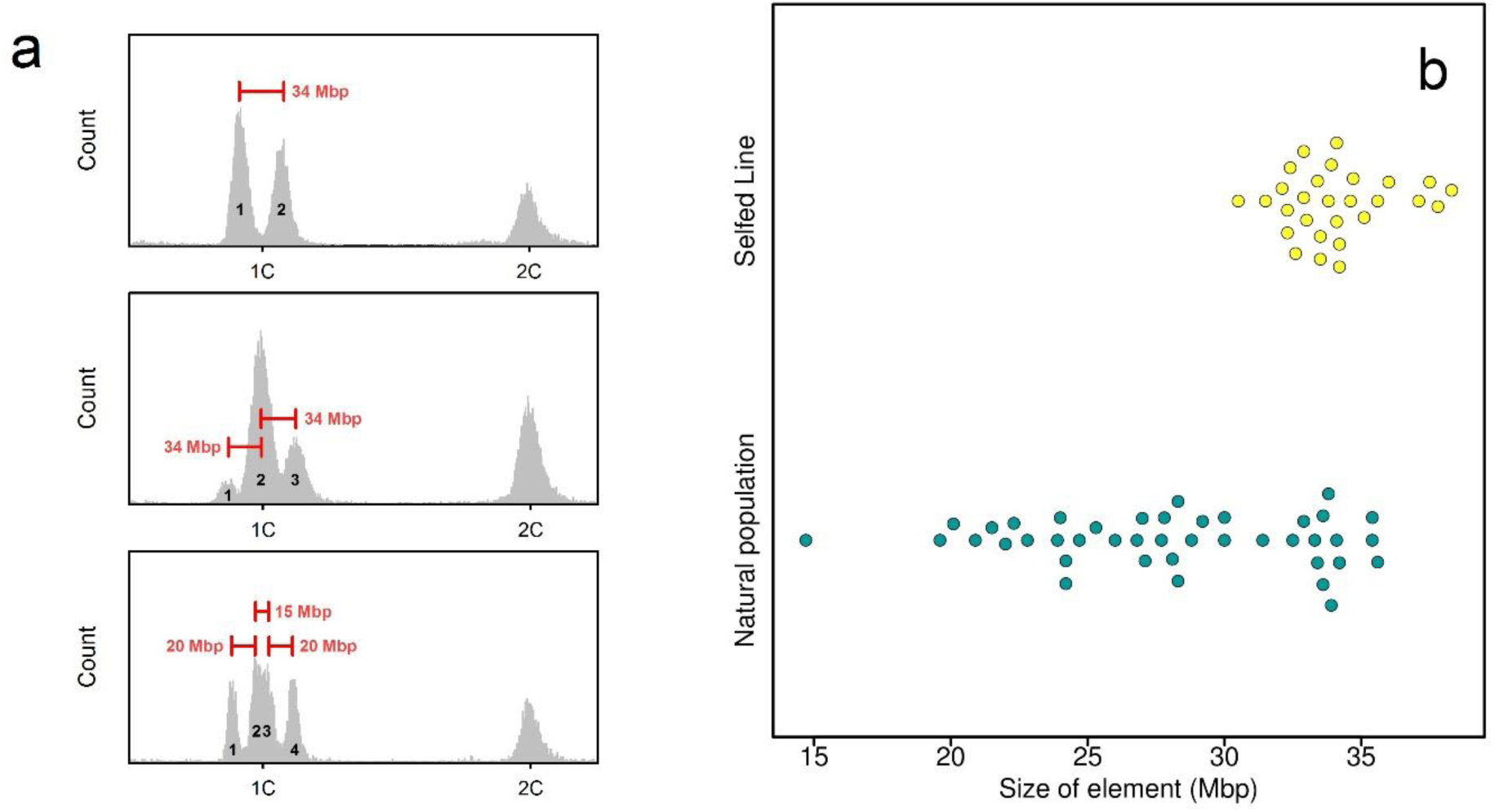
Sizing of genomic elements responsible for male genome size variation. **a** Three examples of size inference, based on the distance between two male peaks (MPs) and knowledge of diploid genome size. Top: Example of a clone with two MPs, which can be explained by segregation of one ∼34Mbp element. Males corresponding to peak 2 contain the 34 Mbp element, while males corresponding to peak 1 are lacking it. Middle: Example of a clone with 3 MPs, which can be explained by two elements of 34Mbp size: Males corresponding to peak 1 are free of elements, males of peak 2 contain one element, and males of peak 3 contain both elements. Bottom: A more complicated case with two, apparently differently sized elements (20Mbp and 35Mbp): Peak 1 corresponds to males without any element, peak 2 to males with the 20Mbp element, peak 3 to males with the 35Mbp element, and peak 4 to males with both elements. **b** Beeswarm plot showing the estimated sizes of elements in the natural population (including their outbred offspring) and in the selfed line (yellow).

### Link between independently segregating elements and genome size variation

The hypothesis of independently segregating elements offers an explanation to many patterns in our data. However, does it provide a *sufficient* explanation for within-population genome size variation? To explore this idea, we formalized our findings in a quantitative model of multiple independently segregating elements (for details and code, see Supplementary Methods). We assumed a minimum diploid genome size at 414 Mbp and that each additional element in a clone would proportionally increase its genome size. Thus, the main input variable of the model is a vector specifying the size and number of elements in a clone. We assumed that all elements segregate completely independently from each other. Thus, if a clone contains, for example, five elements, there is a small chance that it produces males with zero or five elements, while the majority of males will contain two or three elements. In the model, we also defined a parameter for the precision of the flow cytometry measurement, in terms of the coefficient of variance (CV). We set its default value to 2.7%, reflecting the overall mean in our male peak data (Supplementary table 3), but we also explored a CV of 2%, which we obtained in some of our best samples. On the whole, our model can reproduce virtually all male peak patterns found in *B. asplanchnoidis*, and it illustrates some technical issues, e.g., how measurement precision limits the detection of the fine patterns produced by small elements or when a mixture of different element sizes are present in a clone (Supplementary Figs. 9, 10). We could also reproduce the peculiar case of clone OHJ72, our outlier of the natural OHJ-population, with its extremely broad male peak. According to our model, this clone might harbour eleven 34Mbp-elements, or even more, if these elements are smaller.

One method to test whether independently segregating elements provide a sufficient explanation to genome size variation is to “predict” genome size of a clone based on the number and size of its elements. Although this prediction is recursive, since it requires knowledge of female genome size for sizing of the elements, it is not circular, because the distance between male peaks is free to vary. Thus, if we should find that predicted and observed genome sizes differ greatly, this would suggest that independently segregating elements are *not* a sufficient explanation for genome size variation. To explore this idea, we analysed the crossed offspring of the two selfed lines (cf., Fig. 3). This cross was especially suitable because of their low diversity of segregating elements (Fig. 5b, Supplementary table 2). We obtained striking agreement between predicted and observed genome sizes, and could account for 96% of the diploid genome size, on average, just based on the number of 34Mbp elements (Fig. 6a). Nevertheless, all observed genome sizes were approximately 18 Mbp higher than the predicted ones, indicating that there might still be some variation that the model does not account for.

**Fig. 6.**
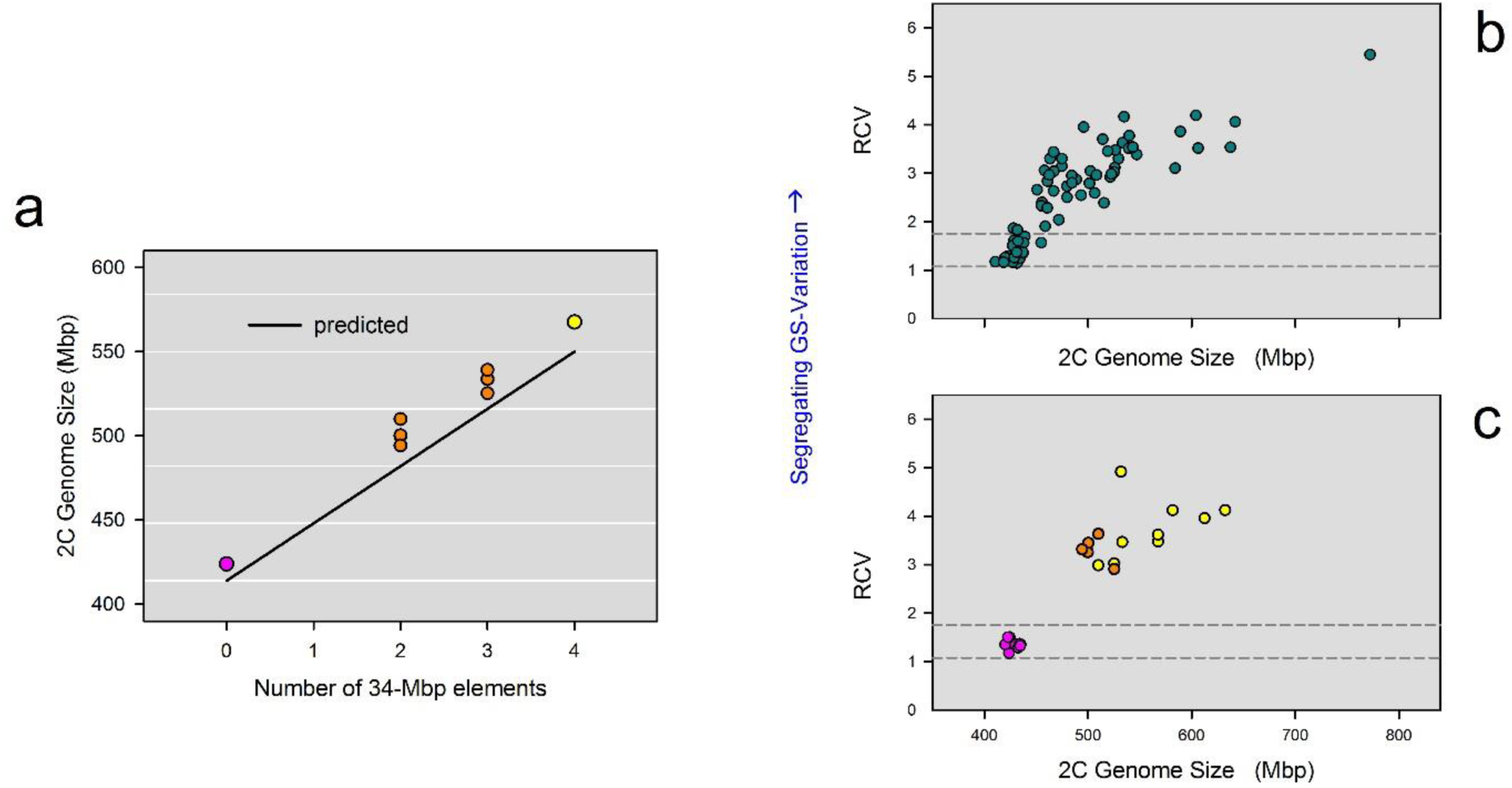
Mechanistic explanations of intraspecific genome size variation.***a*** Prediction of genome size based on the size and number of independently segregating elements, assuming a minimum genome size of 414 Mbp (i.e., free of elements). This figure shows seven clones of the inbred line cross (orange), from which we could determine the exact number of 34Mbp-elements based on their MP patterns. The two parental clones, one with four elements (yellow) and the other with zero elements (purple), are also shown. **b** Genome size *vs*. “relative coefficient of variation” (RCV) in clones of the natural population (Fig. 1a) and their outbred descendants. We used RCV as a proxy for the amount of genomic material that independently segregates during meiosis. RCV was calculated by dividing the CV of all combined male genome sizes (in a clone) by the CV of the female genome size. Horizontal dashed lines are the 95% confidence intervals of the RCV in four outgroup species, which have no intraspecific genome size variation (*B. rotundiformis, B. plicatilis, B. manjavacas*, B. ‘Nevada’). **c** Genome size *vs*. RCV in selfed lines. Selfed line with small genome size in purple, line with large genome size in yellow, and the cross between the two inbred lines in orange.

Identifying the exact size and number of independently segregating elements is not possible for the natural OHJ population as a whole, due to the higher diversity of segregating elements in this population (Fig. 5b), and due to limited resolution of male peak patterns in clones containing multiple differentially sized elements (Supplementary Fig. 9). To circumvent these limitations, we tested whether the amount of meiotically segregating genome size variation is correlated with genome size. To this end, we defined the variable RCV (Relative Coefficient of Variation), which is the CV of all combined male genome sizes of a clone divided by the CV of the female genome size. Thus, RCV accounts for differences in measurement precision across samples (indicated by the CV of the female peak), while it requires only that a clone can produce males in order to be measured, thus extending our analyses to many more clones (n=72). We found that the RCVs of OHJ-clones were considerably higher than those of outgroup rotifer species without genome size variation (Fig. 6b), and that genome size and RCV were positively correlated (Spearman rank correlation, ρ = 0.89, *P* < 0.001). Consistently, RCVs in the “large” selfed line were significantly higher than in the “small” selfed line (Fig. 6c; Mann Whitney U-Test, W = 88, *P* < 0.001). Our model predicted the same curvilinear relationship between genome size and RCV as we observed in *B. asplanchnoidis*, assuming either 20Mbp or 34 Mbp elements (Supplementary Fig. 11). In particular, the prediction based on 20 Mbp elements seemed to match the majority of clones of the OHJ population. Altogether, the positive correlation of RCV with genome size as well as the congruence of the model predictions and the natural OHJ-population suggest that independently segregating elements are largely responsible for genome size variation in *B. asplanchnoidis*.

## Discussion

### Extensive within-population genome size variation in B. asplanchnoidis

Here we report a case of within-species genome size variation in the rotifer *B. asplanchnoidis* and show that even individuals *within* a natural population can differ by 1.33-fold, exhibiting genome sizes ranging from 414 to 552 Mbp. Moreover, artificial selection for large genome size can extend this range to 1.55-fold (640 Mbp) within just two generations. To our knowledge, these ranges represent some of the most pronounced cases of intraspecific variation in animals and plants (Supplementary Table 4).

Claims of intraspecific genome size variation have sometimes met scepticism, due to possible methodological artefacts [35], or the species concept underlying the study [36]. Thus, we used two different internal standards in our flow cytometry measurements, either *Drosophila melanogaster* flies or other rotifer clones that were co-prepared with a focal sample, to demonstrate genome size differences among rotifer clones. Both internal standards yield virtually identical results, confirming that our genome size estimates are accurate and reflect real differences among clones (Supplementary Figs. 2, 3). Previous studies have documented the species status of *B. asplanchnoidis* [32, 37], and have shown that the populations from Austria, Mongolia and Lake Nakuru are genetically distinct, but also experience natural gene flow on a larger geographic scale [32]. Thus, in the case of *B. asplanchnoidis* we can exclude any unrecognized cryptic species.

### Independently segregating elements are responsible for genome size variation

Within-population variation of genome size in *B. asplanchnoidis* is mediated by multiple, large genomic elements that segregate independently during meiosis. These elements range between 15 and 34 Mbp in size and are present in variable numbers in individuals of the natural population. Thus, they account for the full range of genome size variation in this population, including the apparent “outlier” at 792 Mbp. Individuals with the smallest observed genome size are free of such elements, while individuals with larger genomes contain progressively more elements, in different numbers and combinations. Altogether, this allows for a seemingly gradual distribution of genome sizes across the whole population.

The meiotic behaviour of these elements is unusual in that they segregate independently from each other. This has important implications for maintenance of genome size variation in this population, because it facilitates the (re-)generation of a large number of genome size variants in sexual offspring, even if the two parents have identical genome size. For example, consider two parents with three 20 MBp and two 30 Mbp elements, respectively. Both have the same genome size (414 + 60 = 474MBp), but due to independent meiotic segregation of these elements, and recombination, their crossed offspring can have one of ten different genome sizes and range between 414 and 534 Mbp. There is some resemblance to allelic recombination in a genetically controlled trait determined by multiple loci. However, in the case of genome size, variation can even be generated in genetically highly homogeneous lines, such as selfed lines.

On a chromosome level, independently segregating elements in *B. asplanchnoidis* may resemble supernumerary chromosomes (B-chromosomes) that do not pair during meiosis. This is suggested by our results from the selfed line, which seems to have accumulated up to six identical elements derived from an ancestor with three elements. If so, the B-chromosomes are quite large in their relative size, since one 34Mbp element corresponds to 8.2% of the minimum diploid base genome observed in this population. B-chromosomes reported in previous studies were often smaller than normal chromosomes [38], but there are also reports where they reach 3-5% of the diploid genome size [23, 38]. On the other hand, regarding absolute size, B-chomosomes of other species can be twice as large the whole genome size of *B. asplanchnoidis* [23]. A second possible mechanism is that the extra DNA is integrated into the normal set of chromosomes, thus resulting in homeologous chromosomes of different length [e.g., 26]. According to this hypothesis, simple patterns of two or three male peaks could be caused by one or two elements located on different chromosomes, being present only once per chromosome pair. Unfortunately and despite much effort, we could not obtain reliable information on the karyotypes of *B. asplanchnoidis*, which might have allowed us to distinguish between these two possibilities, likely because cell divisions in this organism are limited to a short period when the embryo is encapsulated in a robust eggshell. However, even though we cannot completely rule out the second mechanism, supernumerary chromosomes appear to be the most plausible explanation based on our selfed lines. After all, the two mechanisms are not mutually exclusive. For example in maize, B-chromosomes and “heterochromatic knobs” can both contribute to intraspecific genome size variation [19]. Regardless of whether these independently segregating elements are located on supernumerary chromosomes or integrated in the regular set of chromosomes, we can draw important general conclusions about intrapopulation genome size variation.

### Towards a general model of within-species genome size variation

The model developed here requires as parameters only a minimum genome size (for individuals free of elements), a vector describing size and number of all elements within a clone, and information about their meiotic segregation patterns. This model is consistent with high heritability of genome size and the strong response to selection, while it accounts for the fact that we could not select below a minimum genome size. It is worth mentioning that the minimum genome size that we defined for our population was not obvious from the distribution of the natural population (Fig. 1a), but could only be established by experimental manipulation. Interestingly, previous reports of intraspecific genome size variation in other organisms also report positively skewed distributions (22, 34), which could be an indication of a minimum genome size. Our model could reproduce virtually all observed meiotic segregation patterns (“male peaks”), and it is consistent with the increase of male segregational variation (RCV) with genome size that we observe in the OHJ population (Fig. 6b). Finally, this model can explain the unexpectedly high genome size variation in a selfed line derived from a clone with large genome size.

Our model differs in several notable aspects from previous conceptual models on genome size that were adopted from a perspective of quantitative trait (QT) models. Importantly, QT-models are defined in terms of the mean genome size of a population governed by a large number of insertion and deletion alleles (24). According to our model, QT-models thus refer to the special case where all members of the population have an elevated genome size. However, observations like the minimum genome size at ∼414MBp in the OHJ population, or the fact that genome size variation remained high in the selfed line, suggest that the QT-model might not capture all aspects of the genome size dynamics. In addition, a focus on minimum genome size and DNA-additions, rather than on deletions, is more consistent with the notion that the largest fraction in eukaryotic genomes is excess junk DNA [39]. Nevertheless, it is important to realize that our concept of a ‘minimum genome size’ is not equivalent to a minimal possible genome, but that it is operationally defined, and includes all DNA-additions that are invariably “fixed” in a specific population, or species.

Our present model may be generalized further. Currently, it is strongly oriented towards a mechanism that involves completely independent segregation of extra DNA, which might be best represented in the case of B-chromosomes. However, it can be extended to include other mechanisms, e.g., where extra genomic material is located on normal chromosomes. Accounting for the latter would require some constraints regarding segregation patterns of extra DNA, particularly if these elements are located on the same chromosome. Other variables and parameters, such as minimum genome size and size of elements, would be largely unaffected. Such a generalization has interesting consequences. For example, extra genomic material may become ‘fixed’ in a population once all members carry the same extra DNA element on both chromosomes. Ultimately, a general model would incorporate both mechanisms, integration to normal chromosomes and supernumerary chromosomes, and perhaps even an exchange between both pools [40-43]. Thus, genome size could be highly dynamic on short time scales, due to mechanisms that involve independent segregation, while fixation/loss of extra genomic material on the normal set of chromosomes could explain the long-term changes in genome size across populations or species. Overall, this model does not even contradict long-term gradual (Brownian motion) changes in genome size, while it adds a population-level perspective to taxa that seem to have undergone “saltations” in genome size over macroevolutionary time scales (e.g., [44])

## Conclusions

We have shown that within-population genome size variation in the rotifer *B. asplanchnoidis* is mediated by relatively large genomic elements that segregate independently of each other during meiosis. Our data on short-term artificial selection and inbred line variation suggest that a model that involves only two variables, a minimum genome size and a vector specifying the number, size, and segregation behaviour of the elements that increase genome size is sufficient for capturing most aspects of genome size dynamics in this population. Collectively, our study closes an important gap in our knowledge of how intraspecific genome size variation in populations is mediated by processes at the individual level. Since the genome size variations in this model system are realized across a relatively homogeneous genomic background, this has general implications for identifying the evolutionary forces that are responsible for the immense variation of genome sizes seen across eukaryotes. Most notably, future studies in this system should allow disentangling whether larger genome size can be beneficial, or whether it is always slightly deleterious. In the future, it will also be interesting to elucidate in more detail the mechanisms behind these independently segregating elements, for example, their underlying genomic architecture, or how they interact with the “non-dispensable” parts of the genome.

## Methods

Resting eggs of rotifers were collected in the field from Obere Halbjochlacke (OHJ, N 47°47’11”,E 16°50’31”) and from Runde Lacke (N 47°47’08”, E 16°47’34”), two small alkaline playa lakes in Burgenland (Austria) in 2011. Resting eggs from Lake Nakuru (Kenya) and the two Mongolian clones were obtained from colleagues and have been previously described in detail [30, 32]. All rotifers were cultured as clones, consisting of the asexual descendants of the female that initially hatched from a resting egg (for details, see Supplementary methods).

Genome size measurements were performed with flow cytometry using a detergent-trypsin method and propidium iodide (PI) staining according to [30] with minor modifications (for details, see Supplementary methods). As an internal standard of known genome size, we used the fruit fly, *Drosophila melanogaster* (strain ISO-1, diploid nuclear DNA content: 0.35 pg; [45]).

For sexual crosses between two rotifer clones, we used freshly hatched virgin females and males, which were harvested as eggs from dense rotifer cultures that had initiated sexual reproduction (for details, see Supplementary methods). To analyse inheritance of genome size within a population, we crossed two clones with divergent diploid genome size (414 versus 524 Mbp; called ohj22 and ohj7, respectively) and analysed 27 of their sexual offspring. Two selfed lines were established from the same two clones by growing mass cultures until they produced resting eggs (which were the product of self-fertilization, i.e., males fertilizing females of the same clone). Selfed lines were propagated for three sexual generations by randomly selecting one offspring clone each generation. Finally, we randomly selected one offspring clone from the “large” and “small” line and cross-mated them to produce an inter-line cross.

For the artificial selection experiments, we applied truncation selection to the natural OHJ population by crossing eight clones representing the 10% largest genome sizes among each other (excluding the outlier clone at 792Mbp), and by crossing eight clones representing the 10% smallest genomes, respectively. In the first generation, we used 14 combinations of parental clones for each selection treatment and analysed 1-9 of their offspring. In total, the F1-generation encompassed 31 and 26 offspring clones for the large and small selection-line, respectively. We repeated this selection procedure in the F1-generation to produce a F2-generation. In the small selection line, we used five clone combinations to produce 15 F2-offspring clones. In the large selection line, sexual propensity of the F1-clones was extremely low, limiting us to just one clone combination and 14 of their offspring in F2. The exact genealogy of clones in the F1- and F2-generation can be inferred from their names listed in supplementary table 1. Narrow sense heritabilities *h*^*2*^ were estimated using to the “Breeder’s equation” Δ*Z* = *h*^2^ · *S*, where *S* is the selection differential, and Δ*Z* is the response to selection. Additionally, we calculated *h*^2^ from the slope of the best-fit line for a plot of mid-offspring versus mid-parent genome size of all our available data, which included all crosses and the self-fertilized clones.

To determine the genome size of males relative to (diploid) females, we grew clones from low to high population densities until they started to produce males. We co-prepared males and females from the same clone and subjected them to the same protocol as above, except that we did not use a *Drosophila* internal standard. To better visualize male peaks, we used an excess of males, from 200 males + 100 females to 300 males + 60 females, depending on genome size of the clone. In the flow-cytometry analysis, we quantified the following variables: number of male peaks (up to 6), position of each individual male peak (as the median of the YL-2A value), position of the female peak, standard deviation (YL-2A value) of all combined male peaks, and standard deviation of the female peak. In contrast to the previous genome size measurements, we applied a slightly stricter precision cut-off at 3.5% CV of the diploid female peak, and discarded all samples with higher CVs.

We performed two types of analyses depending on the quality our male peak data. First, in clones that showed multiple discrete male peaks, we counted and sized these peaks (relative to the female peak). We also determined the area under each peak, as a measure for the frequency of each male genome size class. Second, for these and for all remaining clones, we calculated relative coefficient of variation (RCV) according to

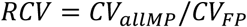

where *CV*_*allMP*_ is the CV across all male peaks and *CV*_*FP*_ is the CV of the female peak. Thus, RCV is a measure for male genome size variation within a sample, corrected for its measurement error (indicated by the CV of the female peak). To obtain a point of reference for RCV values in species without intraspecific genome size variation, we conducted the same analyses in four sister species (*B. rotundiformis, B. plicatilis, B. manjavacas*, B. ‘Nevada’).

## Summary of all supplementary information

### Supplementary information

Supplementary Figures 1–11, Supplementary Table 4, and Supplementary Methods.

### Supplementary file

Supplementary table 1: Genome size estimates of all clones used in this study; 2: Clones used for sizing of independently segregating elements; 3: Clones used for RCV (relative coefficient of variance) calculations.

## Acknowledgements

We thank Alois Herzig for providing access to the sampling sites and help during sampling of Obere Halbjochlacke and Runde Lacke. Sediments from Lake Nakuru were kindly supplied by Steve Omondi and Michael Schagerl. Julie Blommaert helped in preparation of media in some of the experiments. We thank David Mark Welch and Julie Blommaert for helpful comments on the manuscript.

## Authors’ contributions

CPS conceived the study, designed and participated in all experiments. MP and PS contributed the flow-cytometry analysis. SR contributed the line crosses and the selection experiment. AH and MP did most of the culturing work and prepared rotifer biomass for flow cytometry. CPS analysed the data and drafted the manuscript. All authors read and approved the final manuscript.

## Supplementary Information

### Supplementary Figures

**Supplementary figure 1.**
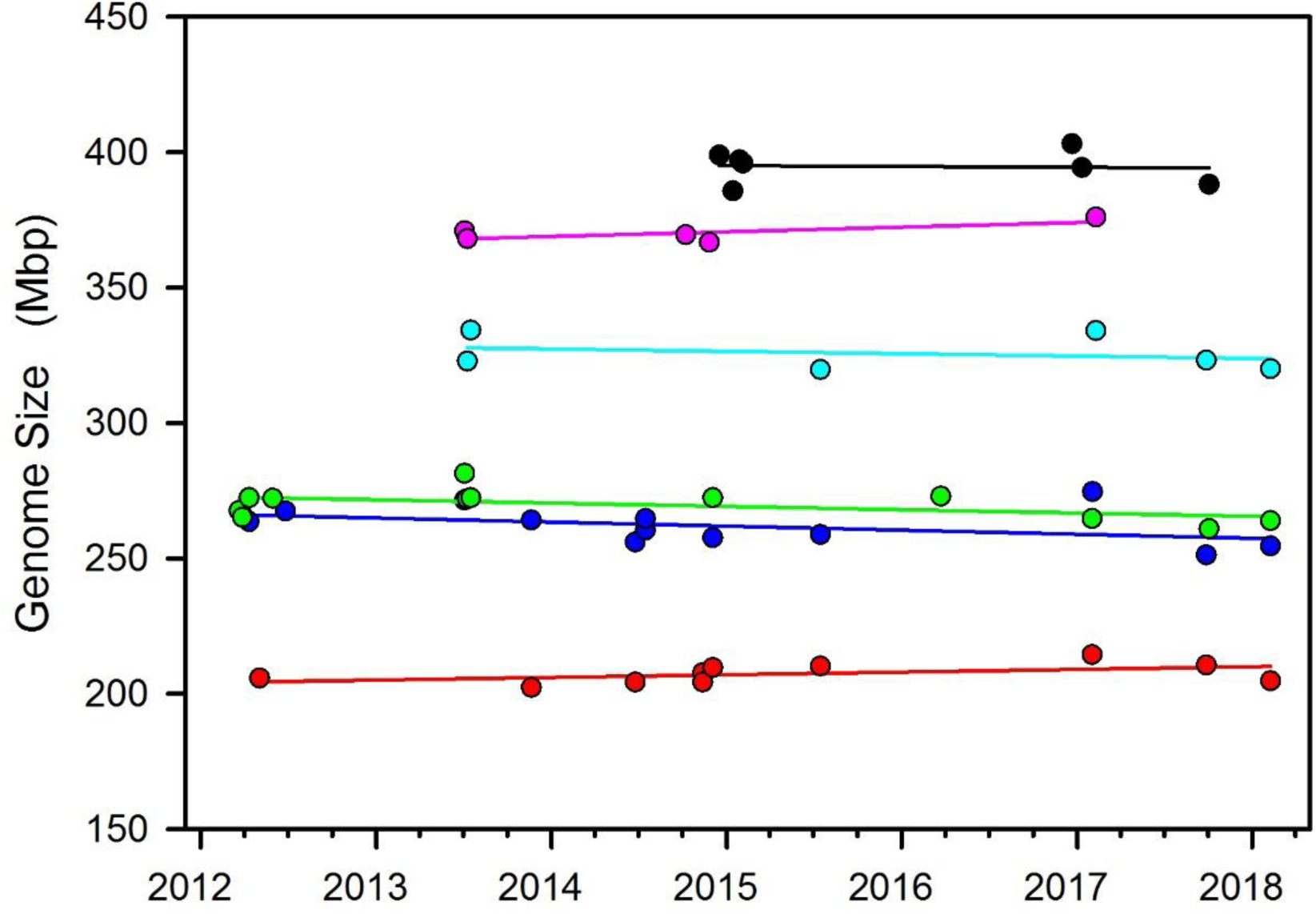
Long-term constancy of genome size during clonal propagation. Time courses of genome size for six *B. asplanchnoidis* clones over a period of > five years (i.e., approximately 600 asexual generations). Colours denote different clones: *ohj22* (red), *ohj7* (blue), *ohj13* (green), *mnchu24* (cyan), *mnchu008* (pink), *ohj72* (black). Each symbol represents an independent genome size measurement.

**Supplementary figure 2.**
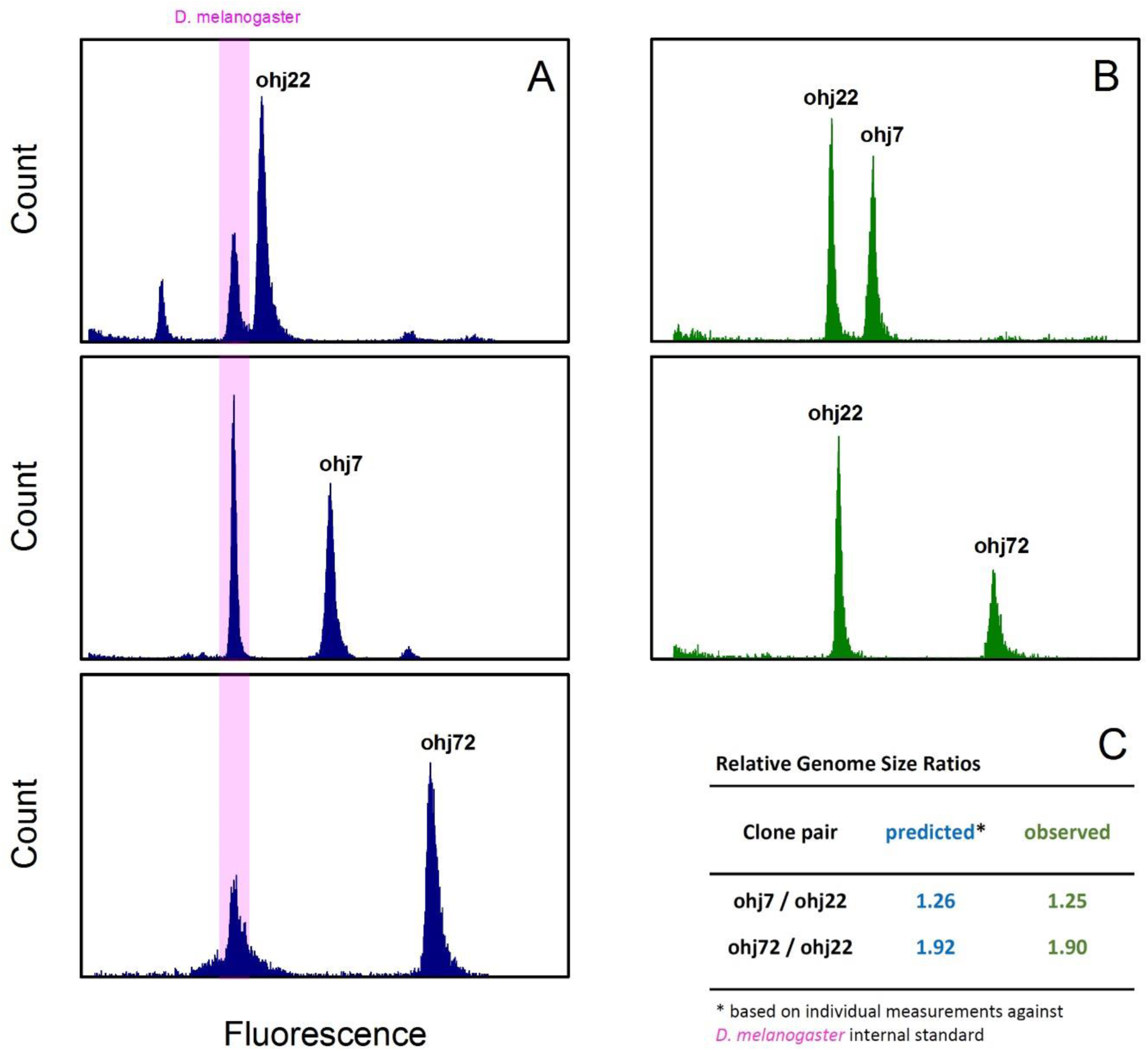
Comparison of two variants of the flow cytometry method for measuring intraspecific genome size variation. **a** Three clones of different genome size were co-prepared with an internal standard of known genome size (*D. melanogaster*, strain ISO 1, 352 Mbp). **b** Pairs of the same three clones were combined into a sample. **C** A comparison of the genome size ratios (= large GS / small GS) determined by the second method shows that these closely matched those predicted by the measurements of the individual rotifer clones, as determined by the first method.

**Supplementary figure 3.**
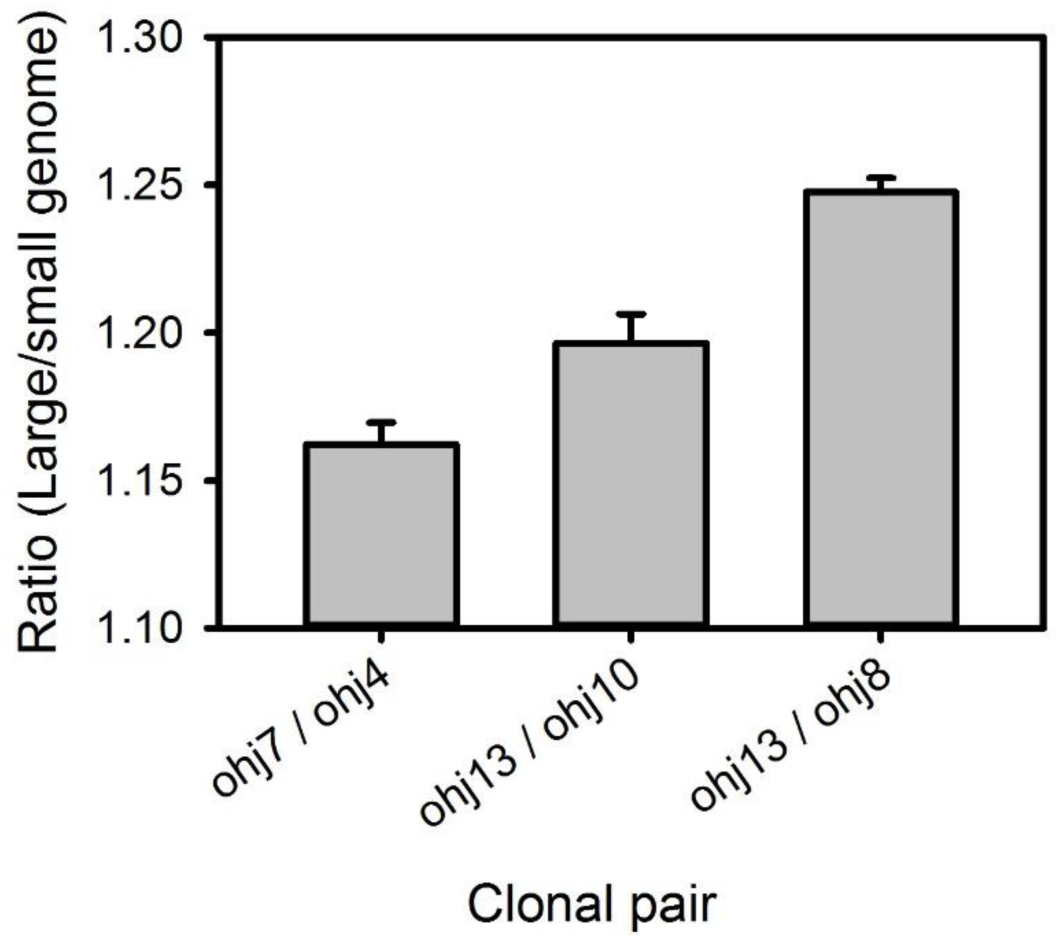
Genome size differences among selected pairs of clones in the OHJ-population. Error bars are standard deviations based on 3-4 replicates (measured on different dates). In each replicate the clonal pairs were co-prepared and stained in the same tube. Kruskal-Wallis test: *H*= 8.95, *d.f*.=2, P=0.011.

**Supplementary figure 4.**
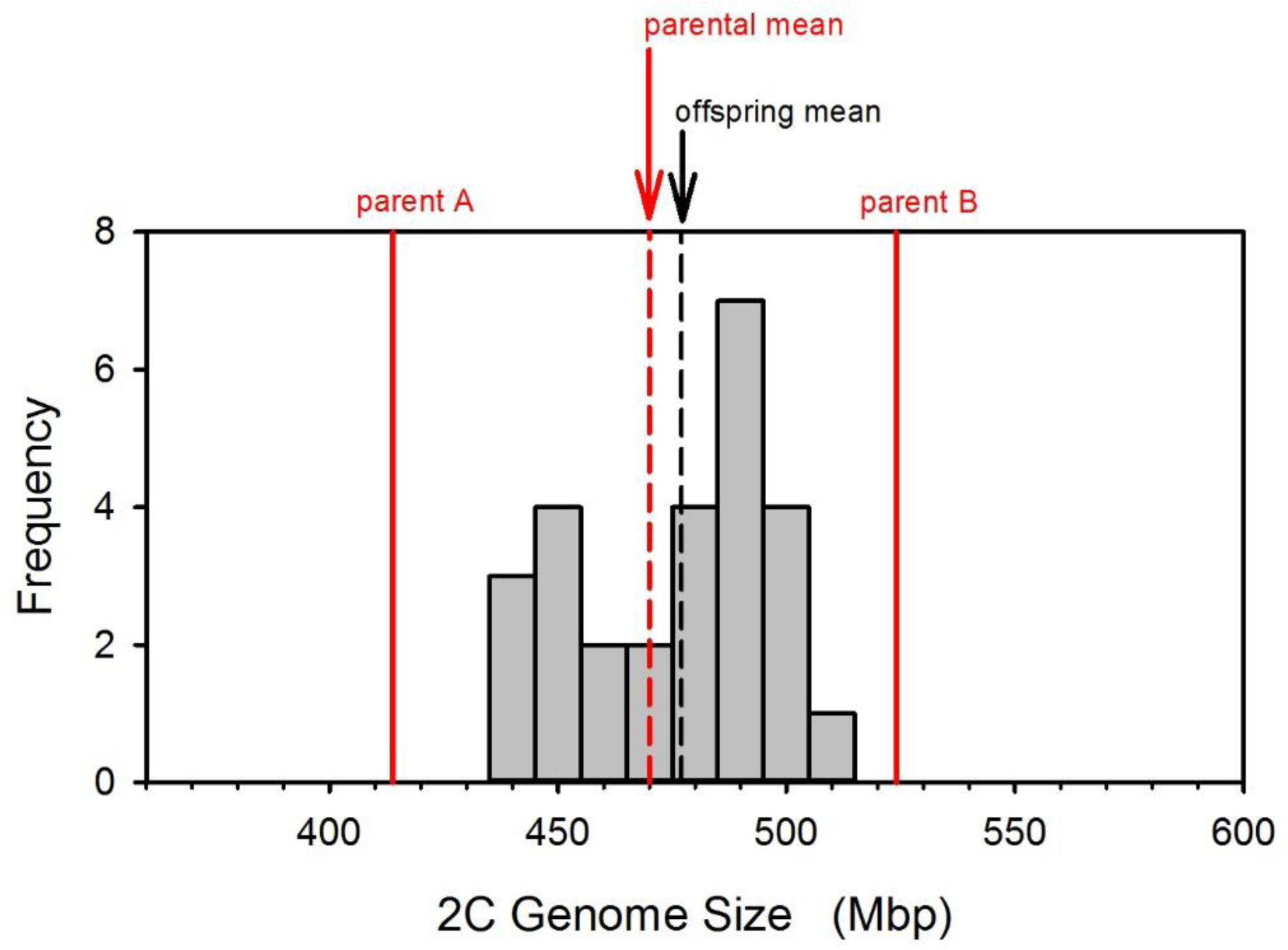
Sexual cross between a clone with large genome size and a clone with small genome size. Parental genome sizes (414 and 524 Mbp) are indicated by the vertical red lines. The dashed red line indicates the parental mean, while the dashed black line indicates the offspring mean genome size. Overall, the 27 offspring genome sizes were intermediate between their parents, even though their distribution was not unimodal.

**Supplementary figure 5.**
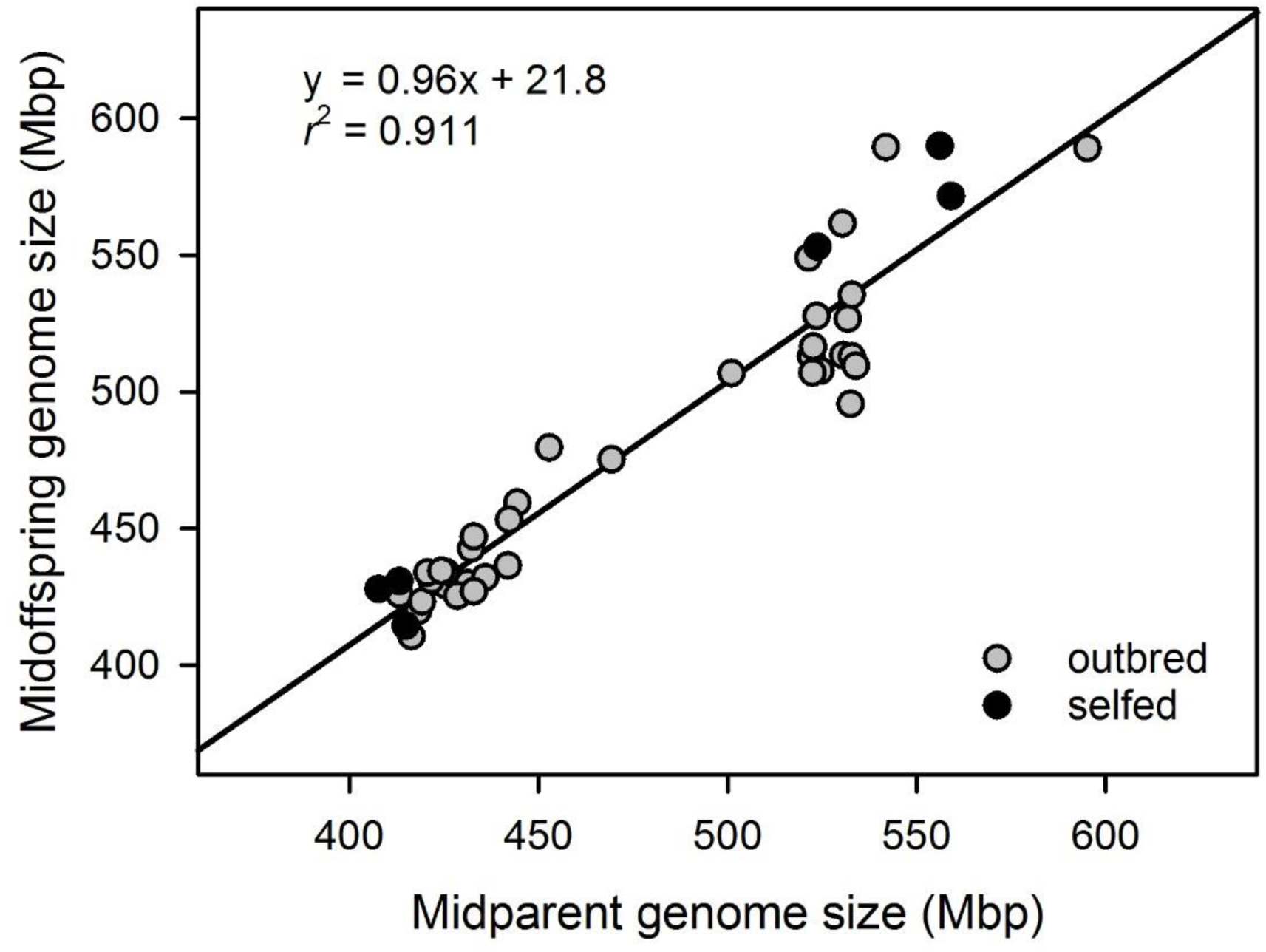
Narrow-sense heritability (*h*^2^) of genome size in *B. asplanchnoidis*. Each symbol represents the mean of parents and offspring genome sizes of a family. This figure combines our results from the artificial selection experiment and crossings between clones with large and small genome size.

**Supplementary figure 6.**
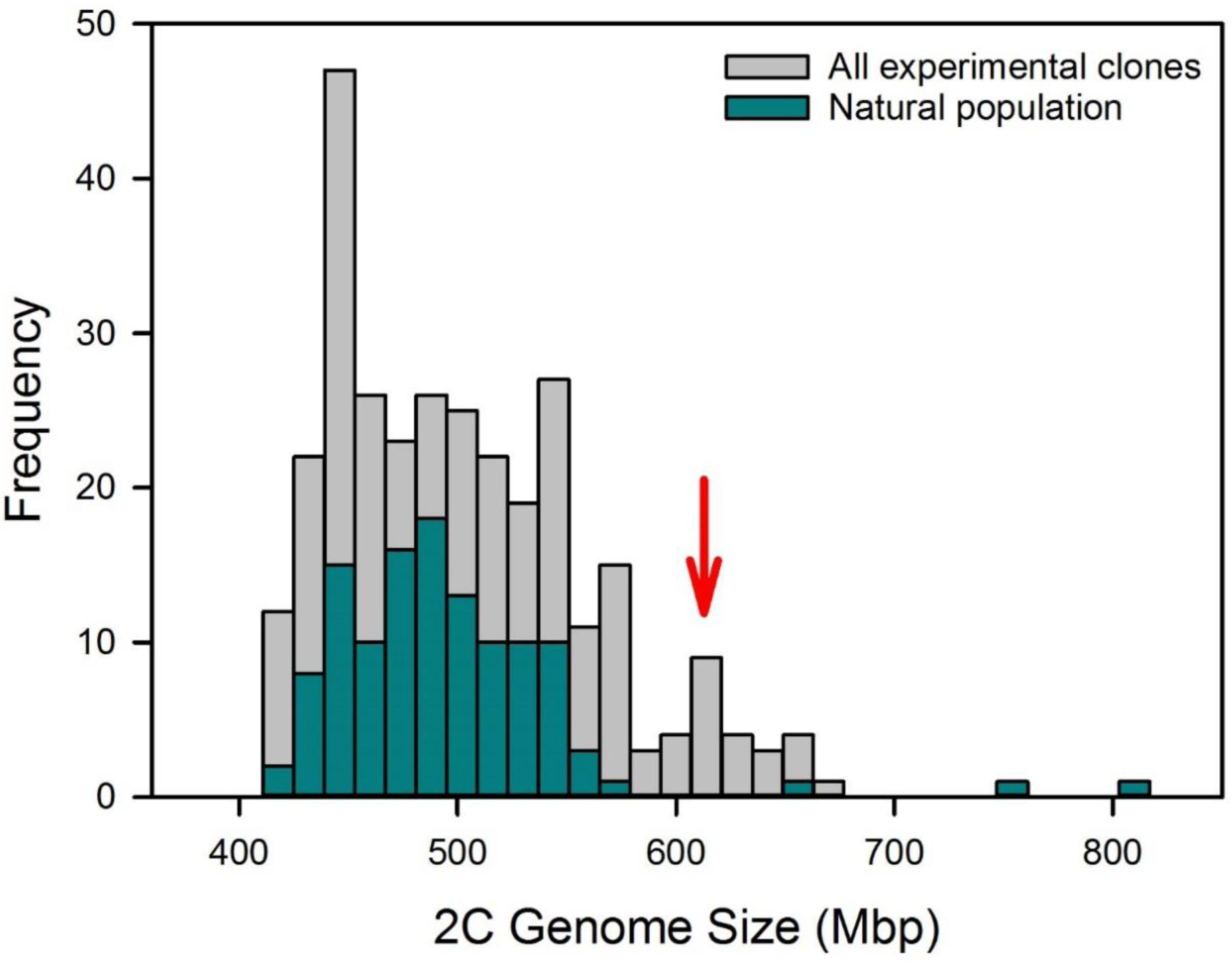
Distribution of genome sizes found in *B. asplanchnoidis*. **Gray**: All *B. asplanchnoidis* clones measured so far (including natural populations, selection lines, and selfed lines). **Green**: Only the natural OHJ population. **Red arrow** highlights genome size classes absent in the natural population, which could be artificially selected in the laboratory.

**Supplementary figure 7.**
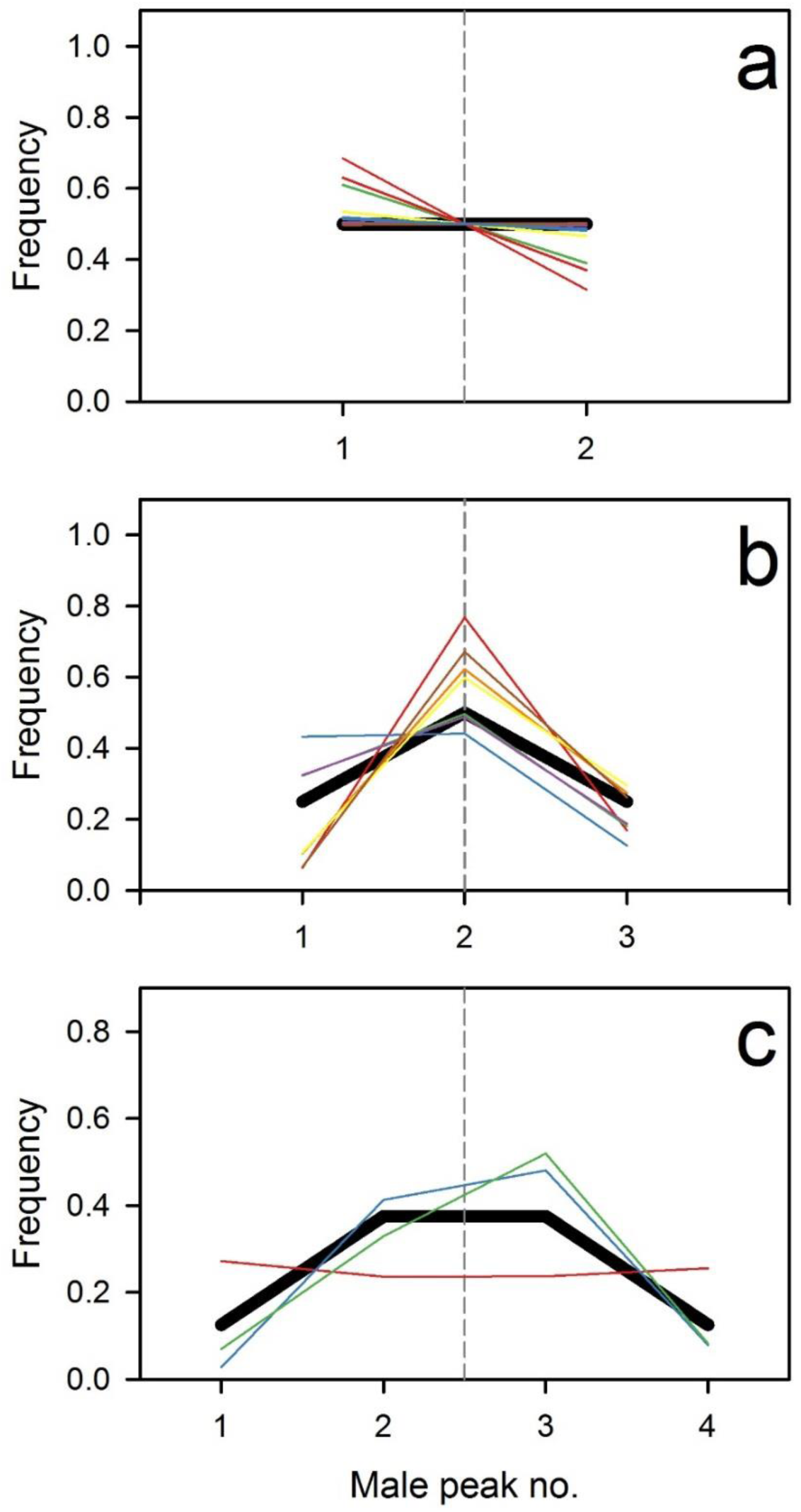
Relative frequency of the different male genome sizes. **a** two malepeaks; prediction (thick line) based on one independently segregating element **b** three malepeaks; prediction based on two equally sized elements, **c** four male peaks; prediction based on three equally sized elements. Coloured lines denote different clones. Dashed vertical line indicates 0.5x diploid genome size.

**Supplementary figure 8.**
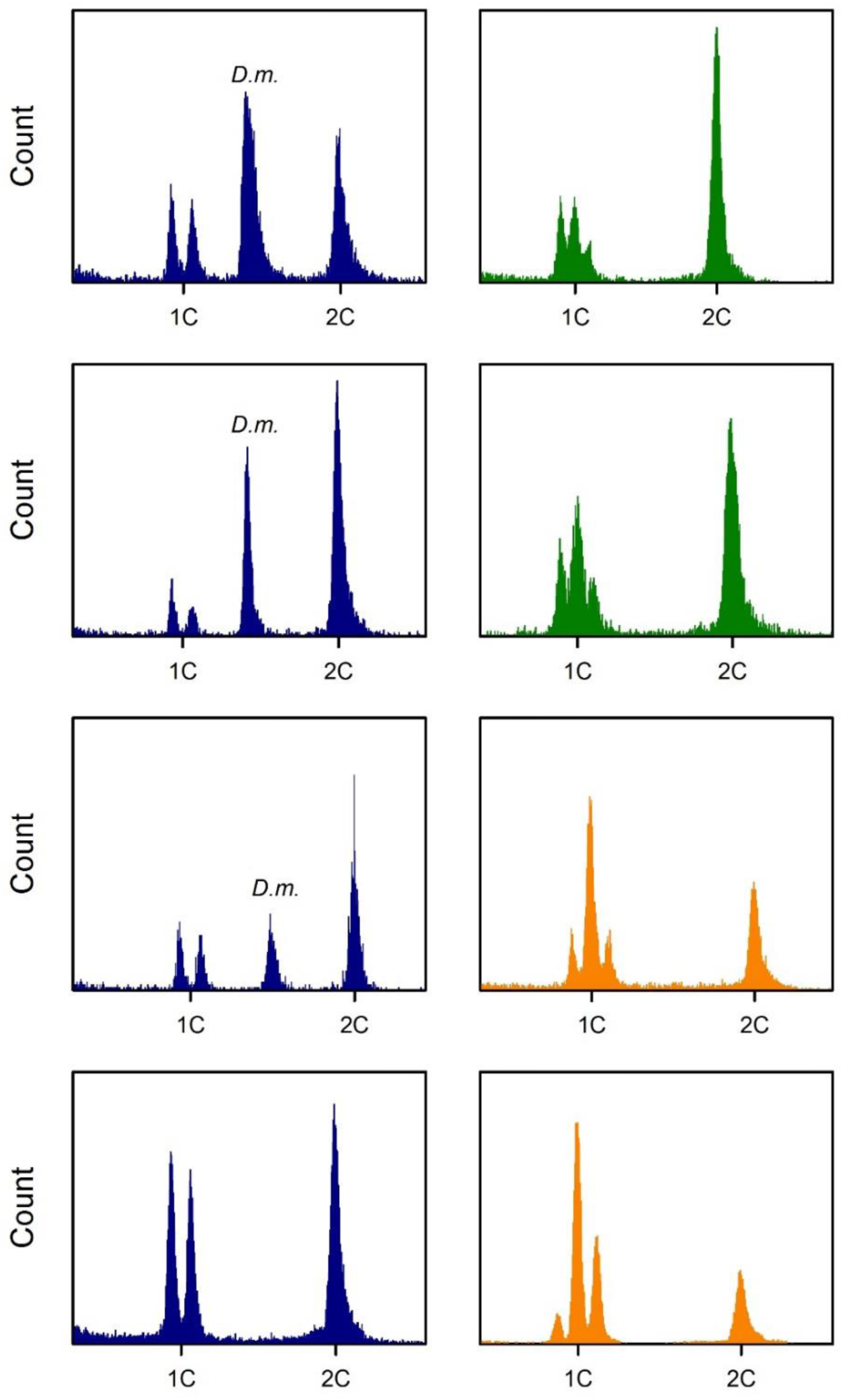
Constancy of “male peak” (MP) patterns in asexually propagated clones. The number of male peaks and their heights (relative to each other) are characteristic for a clone. This figure shows the 2C-female peaks and the corresponding male peaks (peaks scattering around 1C) for three different clones on different dates. **Blue**: Clone *k13×110n2* (two MPs); **Green**: Clone *ohj76* (three MPs); **Orange**: Clone *ohj85* (three MPs). In some of these measurements we used *Drosophila melanogaster* (**D.m.**) as internal standard. Please note that the height of MPs relative to the 2C-female peak might differ among dates, since the number of males per female in a culture is not always the same.

**Supplementary figure 9.**
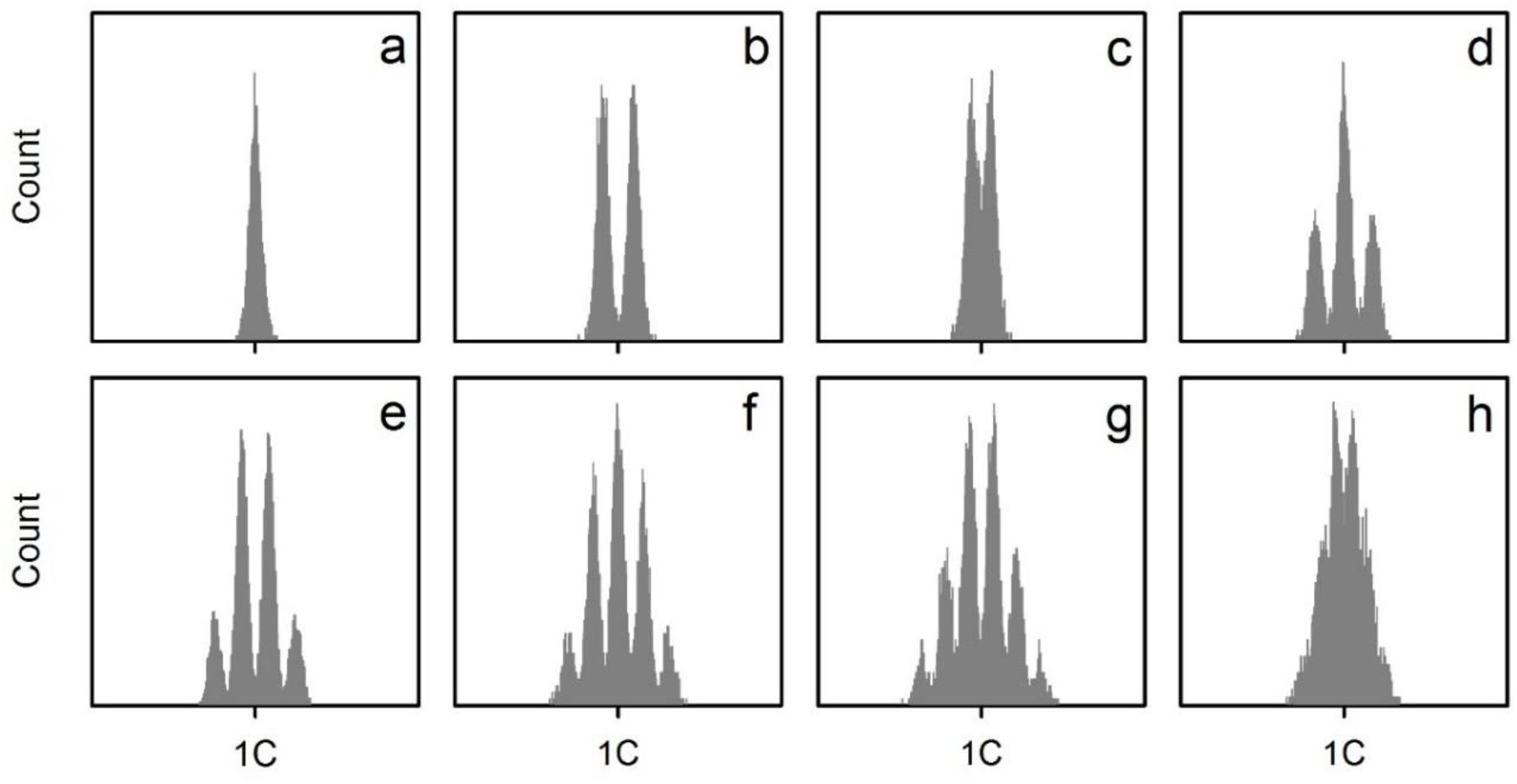
Model simulations of “male peak” (MP) patterns with only one type of independently segregating element. **a** No element; **b** one element of 34 Mbp size; **c** one element of 20 Mbp; **d** two elements of 34 Mbp; **e** three elements of 34 Mbp; **f** four elements of 34 Mbp; **g** five elements of 34 Mbp; **h** five elements of 20 Mbp. The model-parameter ‘measurement precision’ was set to 2.7% (i.e., the coefficient of variance) in all cases.

**Supplementary figure 10.**
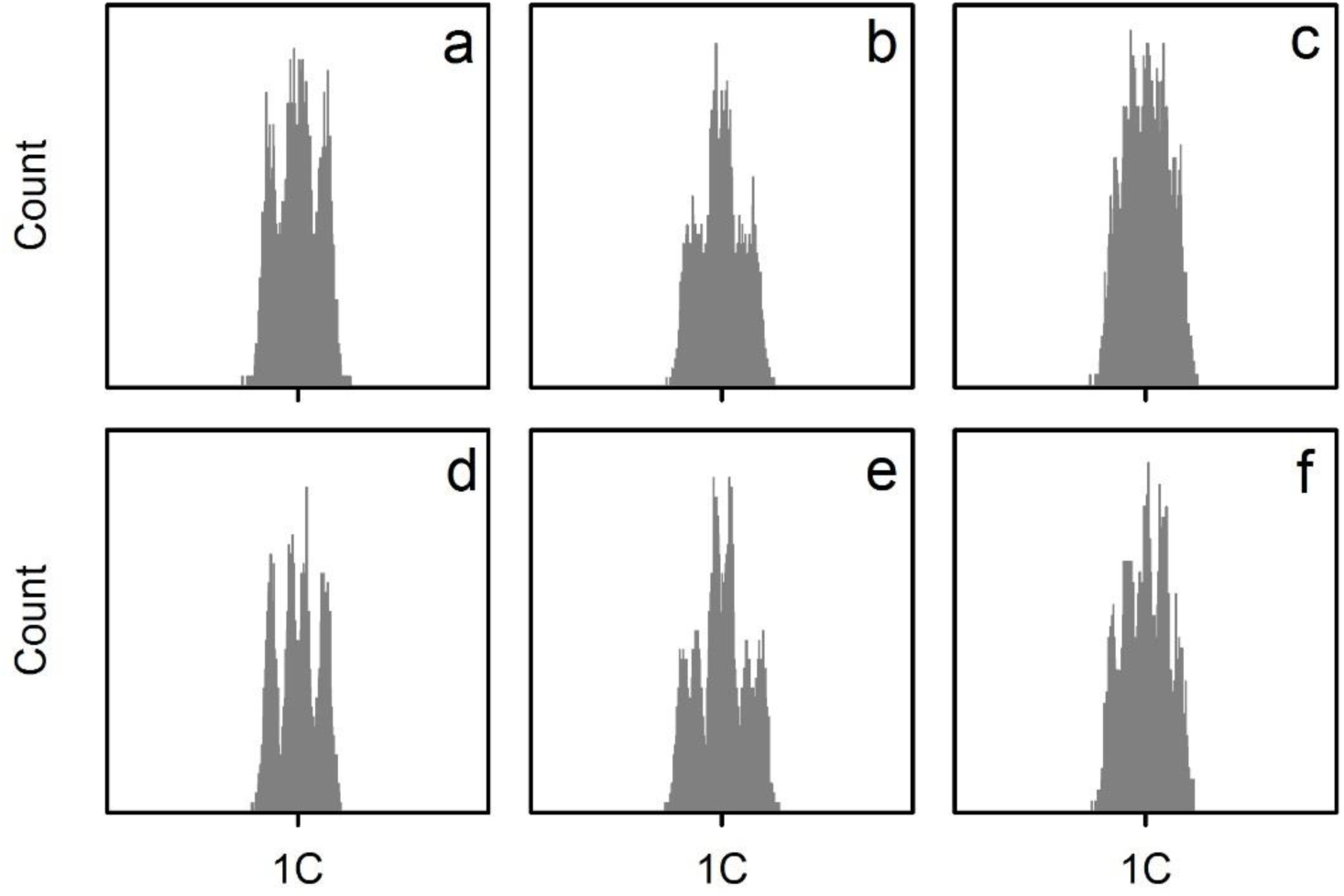
Model simulations of “male peak” (MP) patterns with different types of independently segregating elements. **a** two elements with 20 and 34Mbp; **b** three elements with 15, 34, and 34 Mbp; **c** three elements with 15, 20, and 34 Mbp. The model-parameter measurement precision was set to 2.7% (i.e., the coefficient of variance). **d-f** are identical to **a-c**, except that measurement precision was set to 2.0%.

**Supplementary figure 11.**
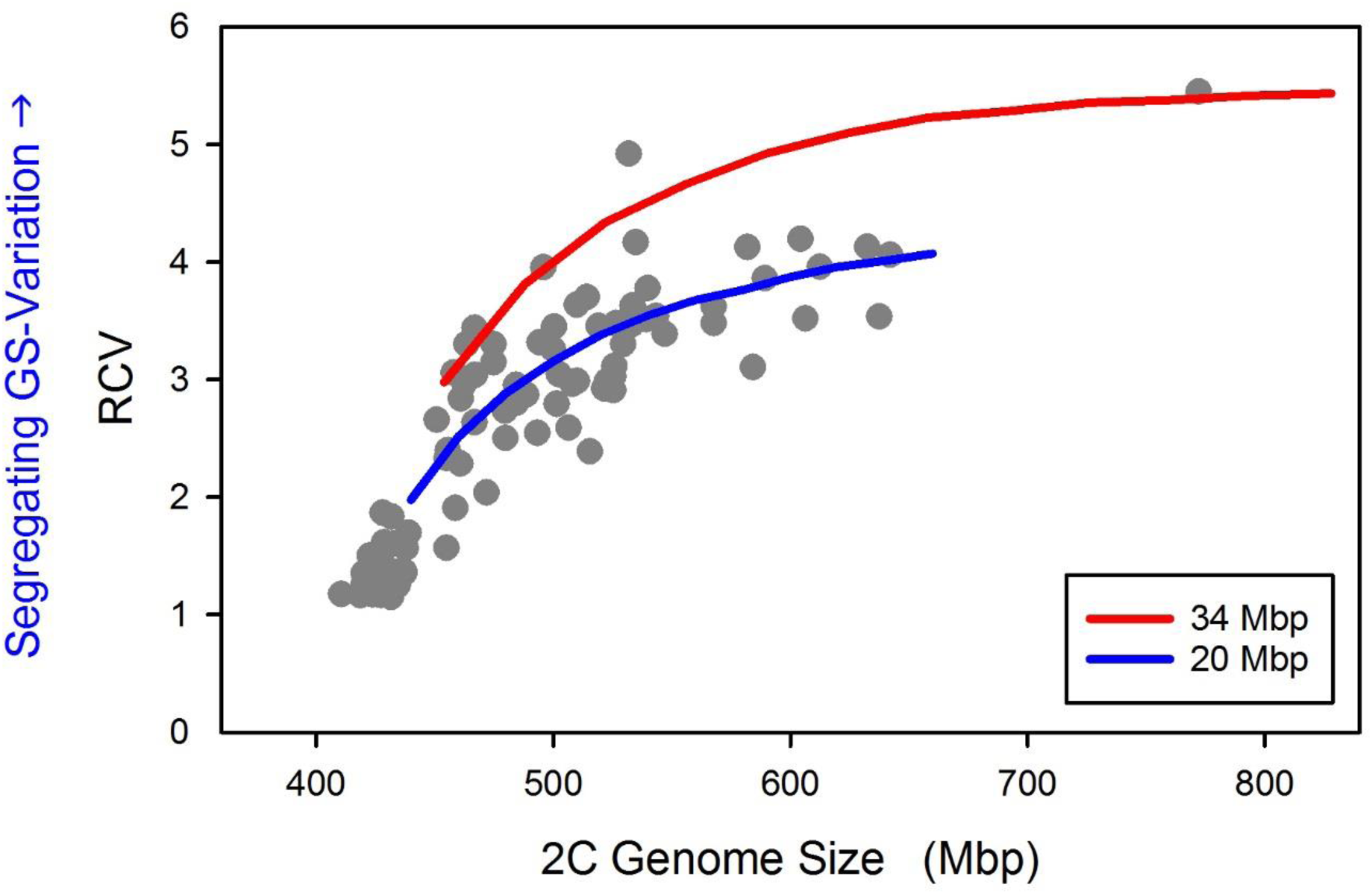
Model simulations of the relationship of RCV and genome size. RCV (“relative coefficient of variation”) was calculated by dividing the CV of all combined male genome sizes (in a clone) by the CV of the female genome size. Red line shows model predictions for genomes with multiple 34 Mbp elements. Blue line shows model predictions for genomes with multiple 20 Mbp elements.. The model-parameter ‘measurement precision’ was set to 2.7%. Dots are all measurements of RCV in this study (cf. Fig. 6 b & c)

### Supplementary Tables

**Supplementary table 4.** Examples of natural intraspecific genome size variation in animals and plants.

| Species name | Common name | 1C-value (pg) | Fold-variation in GS | Reference |
| --- | --- | --- | --- | --- |
| *Zea mays* | Maize | 2.30 | 1.37 | Diez et al. (2013) |
| *Festuca pallens* | Grass | 2.53 | 1.19 | Smarda et al. (2008) |
| *Microseris douglasii* | Coastal silverpuff | 1.20 | 1.20 | Smarda & Bures (2010) |
| *Hordeum spontaneum* | Wild barley | 5.50 | 1.13 | Smarda & Bures (2010) |
| *Arabidopsis thaliana* | Thale cress | 0.16 | 1.14 | Long et al. (2013) |
| *Eyprepocnemis plorans* | Grasshopper | 10.39 | 1.19* | Ruiz-Ruano et al. (2011) |
| *Synalpheus duffyi* | Snapping shrimp | 17.38 | 1.35 | Jeffery et al. (2016) |
| *Drosophila melanogaster* | Fruit fly | 0.18 | 1.15† | Huang et al. (2014) |
| *Brachionus asplanchnoidis* | Rotifer | 0.21 | 1.33 (1.91) | **this study** |

\* assuming individuals with zero vs. six B-chromosomes of 0.64pg size
† variation between inbred strains

### Supplementary Methods

#### Rotifer cultures

Rotifers were cultured in F/2 medium [46] at 16 ppt salinity and with *Tetraselmis suecica* algae as food source (500–1000 cells μl^-1^). Continuous illumination was provided with daylight LED lamps (SunStrip, Econlux) at 30–40 µmol quanta m^-2^ s^-1^ for rotifers, and 200 µmol quanta m^-2^ s^-1^ for algae. During the experiments, rotifers were cultured at 23°C. Stock cultures were kept either at 18 °C, re-inoculated once per week by transferring 20 asexual females to 20ml fresh culture medium, or they were kept for long-term storage at 9°C, replacing approximately 80% of the medium with fresh food suspension every 4 weeks.

#### Genome size measurements

To determine the genome size of rotifer clones, we used flow cytometry and propidium iodide staining of nuclei. Rotifer clones were grown in 250-1000 ml flasks until their population size reached several hundreds to a few thousand individuals. On the day before preparation, rotifers were harvested with 60 µm sieves, and starved overnight in 11 ppt 0.2µm-filtered medium, followed by two washes with filtered medium on the next day. For each sample, we isolated 350 females and subjected them to the flow cytometry protocol. Briefly, rotifers were homogenized in a citrate buffer (3.4 mM Trisodium citrate dihydrate, Nonidet P40 at 0.1% v/v, 1.5 mM Sperminetetrahydrochloride, 0.5 mM Trishydroxymethylaminomethane, pH 7.6), and were then transferred to 750 μl stock solution in a 1 ml Dounce tissue homogenizer. Rotifers were homogenized on ice with 20 strokes using the “tight” pestle of the homogenizer. As internal standard of known genome size we used the fruit fly, *Drosophila melanogaster* (strain ISO-1, nuclear DNA content: 0.35 pg, [45]). After adding two female *Drosophila* heads, the sample was further homogenized with ten strokes. Large debris was removed by filtration through a 40 μm mesh nylon sieve. After addition of 100 μl of 0.021% Trypsin (dissolved in stock solution) the sample was incubated for exactly 10 min at 37°C. To prevent further degradation, 75µl of 0.25% trypsin inhibitor was added (this solution also included 0.05% RNAse A) and the samples were incubated for another 10 min at 37°C. Finally, samples were stained with propidium iodide at a concentration of 50 μg/ml. Stained samples were kept overnight on ice in the dark. Flow cytometric analysis was performed on the next day on an Attune NxT^®^ acoustic focusing cytometer (Thermo Fisher) with an excitation wavelength of 561 nm and a custom-made 590– 650 nm bandpass filter (yellow, YL-2) for detection of propidium iodide fluorescence. Flow cytometric data were analyzed using FlowJo software version 10.0.7r2 (FlowJo LLC). To exclude doublets (i.e., nuclei that pass the detector too close together, thus being recorded as a single “event”) we employed YL2-A *vs*. YL2-H gating. Coefficients of variance (CVs) of individual peaks typically ranged between 1.5% and 4% for both *Drosophila* and rotifers. Very few measurements had CVs higher than 5%, and those replicates were discarded. Conversion from picograms DNA to base pairs were made with the factor: 1 pg =978 Mbp [45]. In most cases, we obtained at least three replicate measurements for a clone, usually on different days and sometimes with several weeks or months apart.

#### Crossings between rotifer clones

For sexual crosses between two rotifer clones, we used freshly hatched virgin females and males, which were harvested as eggs from dense rotifer cultures that had initiated sexual reproduction. Eggs were detached from the females by vigorously vortexing the rotifer culture in 50-ml Falcon tubes for ten minutes. Crossings between clones were accomplished by placing 100 female eggs and 50 male eggs together into the same well of a 24-well plate filled with 750 µl of F/2 medium. After 24 h, when all viable eggs had hatched and animals had time to mate, females were transferred to new wells with fresh food suspension. After a few days, when females started producing eggs, we classified them into asexual females, sexual male-producing females (which were unfertilized), and sexual resting egg producing females. All resting egg producing females were isolated and stored at 7 °C in the dark for at least 2 weeks. To induce hatching, resting eggs were incubated with food suspension at 23 °C and high light intensities (200 µmol quanta m^-2^ s^-1^). Usually after 48 h, the first hatchlings started to emerge, and clonal cultures were initiated.

#### Theoretical model of independently segregating elements

We conceptualized a theoretical model to summarize our finding of independently segregating genomic elements. The main purpose of this model was to test the hypothesis that these genomic elements are *sufficient* as the mechanism of genome size variation. In particular, we examine whether such a model is consistent with our observations of genome size variations in males from clones of different diploid genome size. The basic input parameters and variables of the model are: (i) an assumed minimum diploid genome size of 414 Mbp (for a clone that contains no additional genomic elements), and (ii) a vector describing the size and number of individual elements in a clone (e.g., [34 34 20] for two 34 Mbp and one 20 Mbp element, respectively). The output variable of the model was the predicted diploid genome size of females, calculated as minimum genome size + the summed contribution of all elements. The second output variable was the genome size distribution of haploid males of this same clone (i.e., its “male-peak pattern”). Male genome sizes were calculated as ½ minimum genome size+ the summed contribution of all elements in a particular male. To this end, we assumed that all elements segregate completely independently from each other during meiosis, i.e., each element having an equal chance of ending up in one out of four gametes after the two meiotic divisions. For instance, a clone with two 34 MBp elements should produce 25% gametes/males containing no element, 50% gametes containing one element (50%), and 25% containing two elements. We calculated the male genome size distributions for a total of 10,000 gametes/males per clone. We also simulated measurement errors by assuming a coefficient of variance of 2.7%. This value is based on the average precision of our flow-cytometry measurements of male genome size variation (Supplementary table 3). We also explored CV values of 2%, which we obtained in some of our best samples. The model was written in the MATLAB programming environment (MathWorks®, version R2017A) with the code as follows.

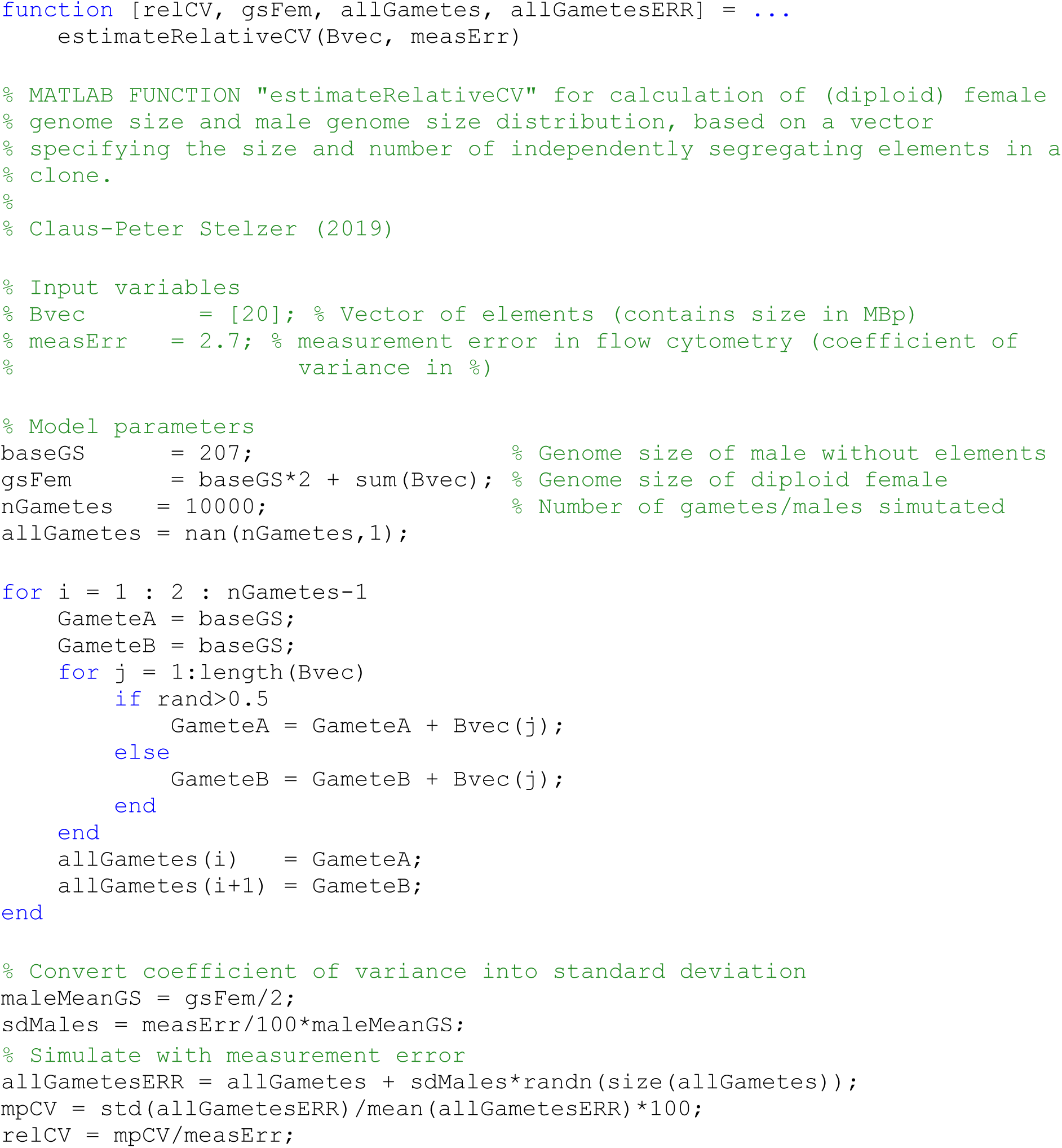

